# Stem cell niche signals Wnt, Hedgehog, and Notch distinctively regulate Drosophila follicle precursor cell differentiation

**DOI:** 10.1101/114090

**Authors:** Wei Dai, Amy Peterson, Thomas Kenney, Denise J. Montell

## Abstract

Adult stem cells commonly give rise to transit-amplifying progenitors, whose progeny differentiate into distinct cell types. Signals within the stem cell niche maintain the undifferentiated state. However it is unclear whether or how niche signals might also coordinate fate decisions within the progenitor pool. Here we use quantitative microscopy to elucidate distinct roles for Wnt, Hedgehog (Hh), and Notch signalling in progenitor development in the Drosophila ovary. Follicle stem cells (FSCs) self-renew and produce precursors whose progeny adopt distinct polar, stalk, and main body cell fates. We show that a steep gradient of Wnt signalling maintains a multipotent state in proximally located progenitor cells by inhibiting expression of the cell fate determinant Eyes Absent (Eya). A shallower gradient of Hh signalling controls the proliferation to differentiation transition. The combination of Notch and Wnt signalling specifies polar cells. These findings reveal a mechanism by which multiple niche signals coordinate cell fate diversification of progenitor cells.

## Introduction

Adult stem cells are important for tissue homeostasis and regeneration due to their ability to both self-renew and generate multiple types of differentiated daughters. Adult stem cells are located in a niche that provides the proper microenvironment to maintain “stemness” ^1,2^. The progeny of stem cells that move away from the niche generally go through a precursor cell (or progenitor cell, transit amplifying cell) stage before they differentiate ^3,4^. However, it is unclear whether the precursor state is simply a loss of stemness due to displacement from niche signals, or whether secreted niche factors might act as morphogens that establish distinct cell fates at different concentrations and distances from the niche.

The Drosophila ovary is an appealing model for studying adult stem cells ^5^. Each ovary contains 16-20 ovarioles, which are chains of egg chambers in increasing stages of maturity ^6^. The anatomy thus displays the temporal sequence of ongoing developmental events (Fig. 1a). Development begins in the germarium, which is located at the anterior tip of the ovariole. The anterior half of the germarium, region 1, contains germline stem cells and their progeny, which continue dividing to produce 16-cell cysts. Somatic escort cells surround the developing cysts as they progress to region 2a. The FSCs are located at the region 2a/2b boundary ^7^, where germ cysts exchange their escort cell covering for the FSC daughters. The posterior half of the germarium contains flattened cysts in region 2b, followed by rounded region 3 cysts. Follicle precursor cells associate with region 2b and region 3 cysts, and their progeny adopt distinct polar, stalk, and main body cell fates, which serve different functions in normal egg chamber development. However the molecular mechanisms that govern these earliest cell fate decisions are unknown and most precursors in region 2b and region 3 do not yet express mature cell fate markers ^8–10^.

**Figure 1.**
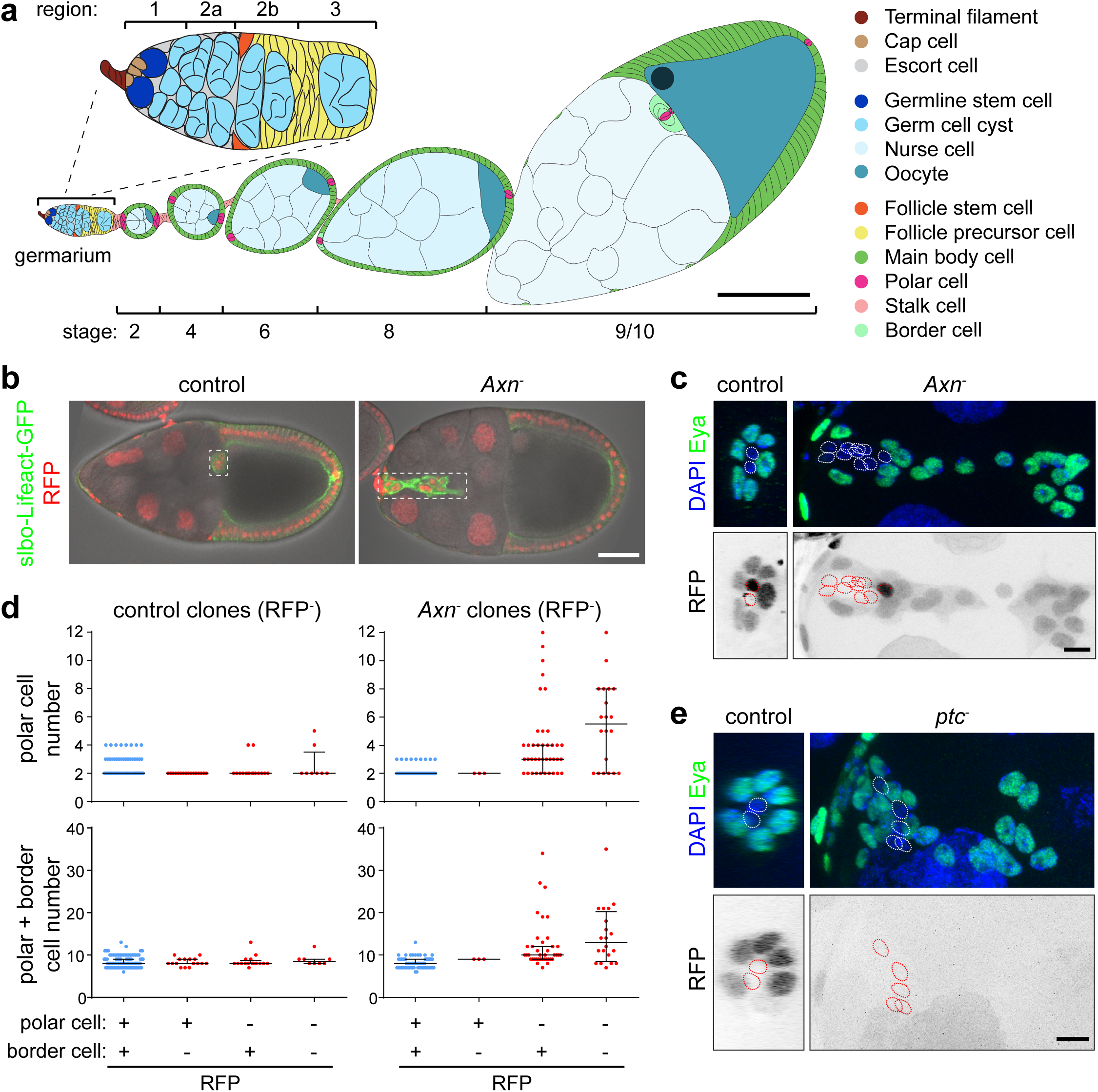
*Axn* mutant clones cause supernumerary polar cells. (**a**) Drawing of a Drosophila ovariole in the sagittal view. Scale bar, 100 μm. (**b**) Sagittal view of stage 10 egg chambers with control (left panel) or FRT82B, *Axn^1511^* mosaic (right panel) border cell clusters (dashed boxes). Scale bar, 50 μm. (**c**) 3D projection view of border cell clusters containing FRT82B control or FRT82B, *Axn^1511^* mosaic clones. Homozygous mutant cells are RFP-negative (RFP^−^). Polar cells are identified by absence of Eya expression (dotted circles). Scale bar, 10 μm. (**d**) Quantification of all border cell clusters in stage 9/10 egg chambers, regardless of whether they have clones or not, in FRT82B control or FRT82B, *Axn^1511^*, 4-5 days after clone induction. Data from n = 284 egg chambers for control, 222 for *Axn^−^*. Each dot represents one border cell cluster. Lines show the median with interquartile range. (**e**) Border cell cluster in FRT42D control or FRT42D, *ptc^S2^* mutant clones. Homozygous mutant cells are RFP^−^. Polar cells are Eya^−^ (dotted circles). Scale bar, 10 μm.

Several signalling pathways have been implicated in regulating follicle precursor cell fate specification and differentiation. Notch signalling is required for polar cell specification ^9^ and is present in mature polar cells at high levels in stage 1 ^11^. Earlier Notch activity at the region 2a/2b boundary is required for the migration of one FSC daughter laterally across the germarium, while other daughters move posteriorly ^8^. However, Notch activity does not seem to be sufficient to induce ectopic polar cells in the main body region ^10^, raising the question whether additional factors are required for polar cell specification. Follicle stem and precursor cells receive signals secreted from the niche escort cells, in addition to germline Delta, which activates the Notch receptor on follicle cells. Niche factors including Wnt, Hh, epidermal growth factor, and bone morphogenetic protein, which are crucial in many adult stem cell niches, are important for FSC maintenance ^12–18^. Hyper-activation of Wnt or Hh signalling causes defects in follicle cell differentiation ^12,19^, but the origin of these phenotypes is not understood and it remains unclear whether these niche signals normally regulate progenitor cell fate or differentiation.

In a forward genetic screen for mutations that disrupt cell fates in the ovary, we identified a mutant allele of *Axin* (*Axn*), a negative regulator in the Wnt pathway. Superficially, the phenotype resembled that caused by mutations in *patched* (*ptc*) or *costal* (*cos*), two negative regulators of Hh signalling. However, we traced both defects back to the earliest steps of follicle cell specification. We developed quantitative analyses of the differentiation markers, Eya and Cas, as well as Wnt, Hh and Notch signalling reporters to reveal distinct roles of these three pathways. We found that a Wnt signal from the FSC niche maintains proximally located progenitor cells in a multipotent state by suppressing the cell fate determinant Eya, whereas a graded Hh signal delays differentiation of all follicle cell types. Notch coordinates with Wnt and Hh in both space and time to specify polar cells. The combination of these three signals produces appropriate spatial patterning of cell types and temporal patterning of differentiation.

## Results

### Distinct effects of Wnt and Hh hyper-activation on follicle cell differentiation

At stage 8 of oogenesis, anterior polar cells specify neighbouring epithelial follicle cells as motile border cells, and together they migrate as a cluster during stage 9 (Fig. 1a). In a forward genetic screen of EMS-induced mutations that cause border cell defects in mosaic clones ^20^, we identified a line that produced abnormally large border cell clusters. Compared to control clusters, which are usually composed of 5-7 migratory cells surrounding two polar cells, clusters containing mutant cells showed as many as 6-12 polar cells and 14-35 total cells per cluster (Fig. 1b-d). The phenotype was autonomous to the polar cells, as supernumerary polar cells and over-sized border cell clusters were only observed when polar cells were homozygous for the mutation (Fig. 1d). The supernumerary polar cell phenotype resembled those previously reported for *eya* ^21^, *cos* ^22^, and *ptc* (Fig. 1e; ^19^). We mapped the new mutation to genomic location 99D3 (Supplementary Fig. 1a-b), which contains the *Axn* gene. *Axn* allele S044230 produced a similar phenotype (Supplementary Fig. 2) and failed to complement the new mutation for lethality. We therefore named the new allele *Axn^1511^*. This supernumerary polar cell phenotype was somewhat surprising as it had not previously been reported for the *Axn* gene.

In addition to supernumerary polar cells, *Axn^−^* and *ptc^−^* clones showed abnormal stalks, consistent with previous reports ^12,23^. Polar and stalk cell specification occurs early in the germarium, and these cells stop dividing soon after they exit the germarium ^7,24^. In the ovary, hyperactive Wnt or Hh signalling affects differentiation ^12,19^. We asked exactly what aspects of differentiation were affected by Wnt or Hh hyper-activation.

We used two complementary markers, Eya ^21^ and Cas ^25^ (Fig. 2a). To be comprehensive, we performed 3D reconstructions of ovarioles and quantified the levels of both Eya and Cas in every somatic cell (Fig. 2b; Supplementary Fig. 3). We found barely detectable levels of either protein in regions 1 and 2a escort cells. Low but increasing levels of both Eya and Cas were present in FSCs and their immediate daughters in region 2b^A^ and 2b^P^. Eya and Cas show differential expression in region 3/stage 1, such that some cells expressed higher levels of one or the other (indicated by divergence from the diagonal in the graphs in Fig. 2b). As development proceeded through stages 1-4, the levels of Eya and Cas diverged more and more. By stage 4, Eya and Cas became completely distinct markers for main body (Eya^+^ Cas^−^) versus polar/stalk (Eya^−^ Cas^+^) fates (Fig. 2a-b). Thus, Eya and Cas are excellent markers for studying the earliest cell fate diversification in the ovary.

**Figure 2.**
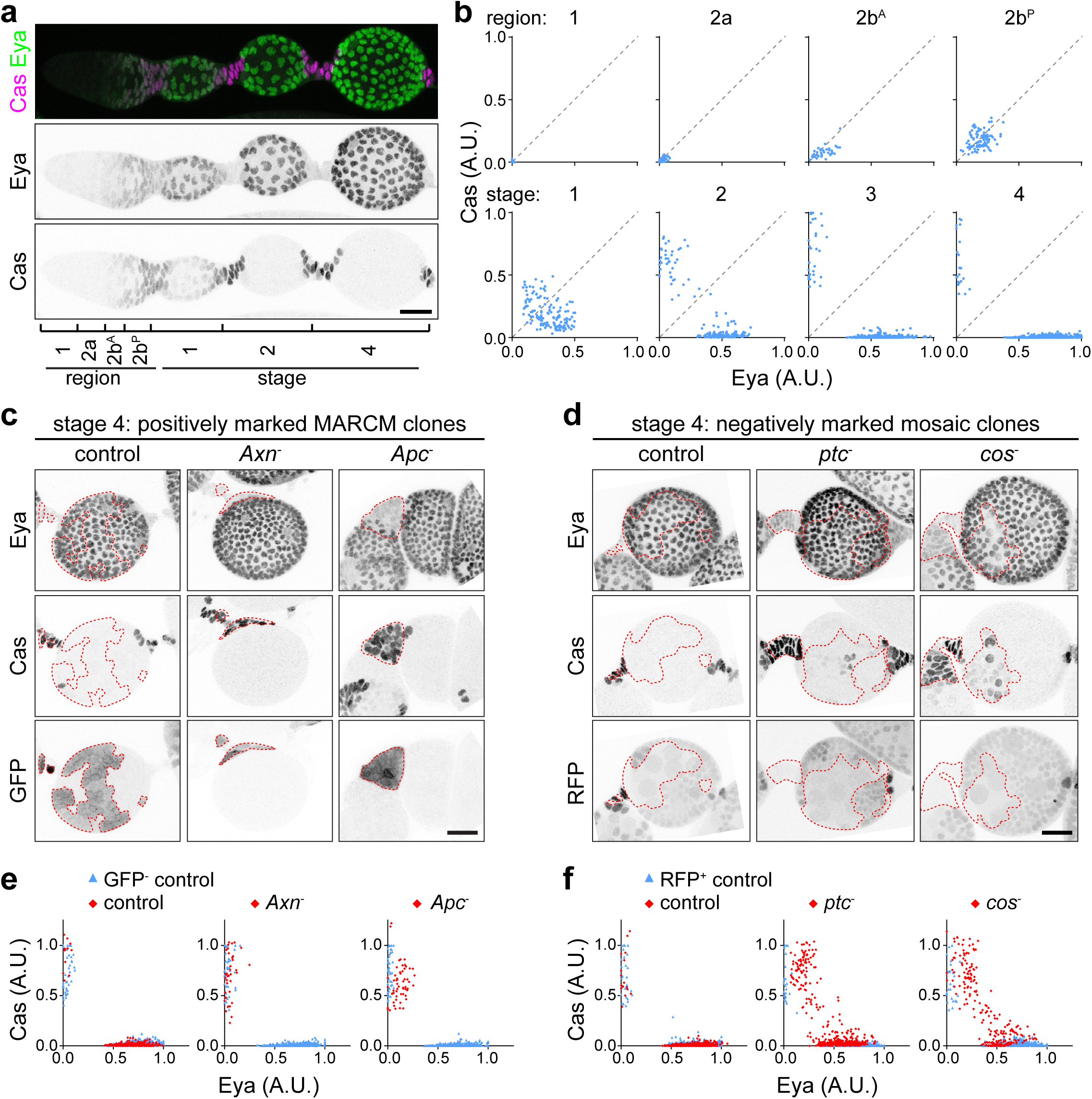
Differential effects of Wnt and Hh hyper-activation on follicle cell differentiation. (**a**) 3D projection view of one half of a wild type ovariole from the germarium to stage 4, stained with Eya (green) and Cas (magenta) antibodies. Individual channels are shown in black and white. (**b**) Quantification of Eya and Cas fluorescence intensity in all somatic cells in germarium regions 1-3 and in stage 1-4 egg chambers. Data from n = 2,122 cells from 3 ovarioles per stage. Each dot represents one cell. Data were normalized to maximum Eya or Cas intensity per ovariole. (**c**) Stage 4 egg chambers with FRT82B control mosaic FSC clones compared to FRT82B, *Axn^1511^* or FRT82B, *Apc2^g10^, Apc*^Q8^. Homozygous mutant cells in the main body and anterior stalk regions are outlined (GFP^+^, dashed lines). Eya+ GFP^−^ cells appear in the outlined *Apc^−^* clone due to Z stack projection. (**d**) Stage 4 egg chambers with FRT42D control, FRT42D, *ptC^S2^* or FRT42D, *cos^H29^* mosaic FSC clones. Homozygous mutant cells are RFP^−^ (dashed lines). (**e-f**) Quantification of Eya and Cas fluorescence intensity in mosaic FSC clones. Data from n = 1,222-1,447 cells from 4 stage 4 egg chambers per genotype. Data were normalized to maximum Eya or Cas intensity in control cells per egg chamber. Scale bars, 20 μm. A.U., arbitrary unit.

To examine the differentiation defect caused by hyperactive Wnt or Hh signalling, we made FSC clones and stained them for Eya and Cas (Fig. 2c-d). Hyperactive Wnt signalling by loss of the β-catenin [or Armadillo (Arm) in Drosophila] destruction complex component *Axn*^−^ or *adenomatous polyposis coli* (*Apc^−^*) produced many egg chambers containing only Eya^−^ Cas^+^, polar/stalk-like mutant cells, in contrast to control clones in which Eya^+^ cells were frequent (Fig. 2c,e). In contrast, FSC clone with hyperactive Hh signalling by loss of the negative regulators *ptc^−^* or *cos*^−^ were not biased toward polar/stalk fates.

They instead produced many Eya^+^ Cas^+^ cells in stage 4 that resembled control cells in stage 1-2. Eya^+^ Cas^+^ cells were virtually never observed in controls in stage 4 (Fig. 2b,d,f). These results suggested distinct differentiation problems in Wnt or Hh hyper-activation, and prompted us to examine their cause and the endogenous function of Wnt and Hh in further detail.

### Wnt and Hh act independently in the germarium

Wnt and Hh signalling positively regulate one another in some settings ^26^, while they antagonize ^27,28^ or play independent roles in other cases ^29^. In the ovary, Wnt and Hh are stem cell niche factors produced in cap cells and escort cells ^16–18, 30^. To understand their relationship, we examined Wnt and Hh activity patterns. We used *frizzled 3* (*fz3*)*-RFP* ^31,32^, which is a reporter for Wnt signalling activity, in order to assess the pattern of Wnt pathway activation. *fz3-RFP* was highly expressed in region 1-2a and showed a graded pattern in region 2b (Fig. 3a-b). The *fz3-RFP* signal was reduced in *armRNAi* expressing clones and increased in *Axn^−^* clones, demonstrating that it is indeed responsive to Wnt signalling (Fig. 3c-d; Supplementary Fig. 4a). *ptc-GFP* is a reporter for Hh signalling ^18^ and shows a pattern similar to *fz3-RFP* (Fig. 3e-f), though the gradient appears to extend more posteriorly. The *ptc-GFP* signal was reduced when knocking down *smoothened* (*smo*), a positive regulator in the Hh pathway (Fig. 3g-h; Supplementary Fig. 4b). Unexpectedly, *cosRNAi* also caused a reduction of *ptc-GFP* signal in region 2b (Fig. 3g-h; Supplementary Fig. 4c), while in later stages the signal increased as expected for loss of a negative regulator (Supplementary Fig. 5). To decipher the relationship between Wnt and Hh signalling in the germarium, we examined the pattern of *fz3-RFP* in *smo*^−^ or *ptc*^−^ clones and the pattern of *ptc-GFP* in *dishevelled* (*dsh*, a positive regulator in the Wnt pathway) or *Axn* mutant clones. Changing Hh signalling had no detectable effect on the Wnt activity pattern in region 2b (Fig. 3c-d), nor did changing Wnt signalling influence Hh activity (Fig. 3g-h). Thus, Wnt and Hh appear to function independently in the ovary.

**Figure 3.**
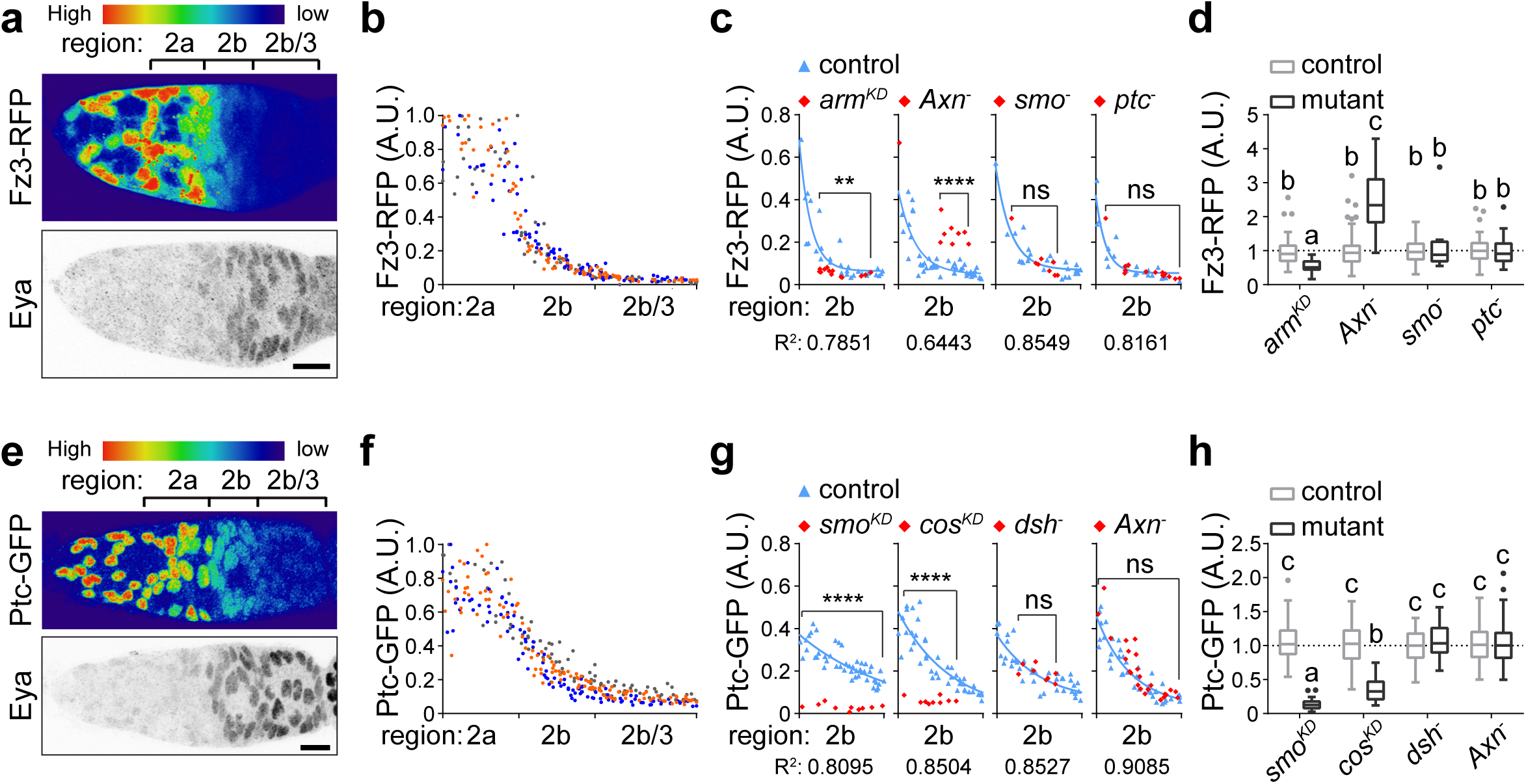
Independent actions of Wnt and Hh. (**a**) 3D projection view of one half of a germarium expressing the Wnt activity reporter *fz3-RFP* and stained for Eya. RFP intensity is displayed using the “Physics” lookup table. (**b**) Quantification of Wnt reporter intensity in all somatic cells from region 2-3 along the anterior-posterior axis. Data from n = 276 cells from 3 germaria. Each dot represents one cell. The three colours represent the three different germaria. (**c**) Wnt reporter activity in germaria with *armRNAi, Axn^S044230^, smo^D16^*, or *ptc^S2^* mutant clones (red diamonds) compared to control cells in the same germarium (blue triangles). R^2^ quantifies goodness of fit of nonlinear regression. (**d**) Quantification of Wnt reporter activity in germaria with mosaic clones. Data (median with interquartile range) from n = 16-103 cells from 3 germaria per genotype. Data were normalized to the predicted value on the one phase decay curve fitted on the internal control cells. (**e**) A germarium expressing the Hh activity reporter *ptc-GFP*. (**f**) Quantification of Hh reporter intensity in all somatic cells from region 2 - 3 along the anterior-posterior axis. Data from n = 314 cells from 3 germaria. (**g**) Hh reporter activity in germaria with *smoRNAi, cosRNAi, dsh*^***3***^, or *Axn*^***S044230***^ mutant clones. (**h**) Quantification of Hh reporter activity in germarium with mosaic clones. Data (median with interquartile range) from n = 18-105 cells from 3 germaria per genotype. Scale bars, 10 μm. A.U., arbitrary unit. ^*\*\*,p*^ < 0.01; ^****^, *p* < 0.0001; Samples labelled with different letters are significantly different at *p* < 0.01.

### Wnt signalling inhibits expression of the main body fate determinant Eya

*Axn^−^* FSC clones frequently gave rise exclusively to Cas^+^ cells (Fig. 4a-b). *Axn^−^* clones appear in the normal polar/stalk region, or as small clones in the main body region forming ectopic polar and stalk cells, or as large clones that form a continuous stalk with a single polar cell cluster, causing the egg chamber to appear to bud from the side. Clones generated at a later stage, however, did not show this cell fate bias (Supplementary Fig. 6; ^12^), suggesting a narrow developmental time window for Wnt signalling to affect cell fate. To understand how hyper-activation of Wnt created clones biased toward Cas^+^ polar/stalk-like fate, we considered a few possibilities. The *Axn^−^* Eya^+^ main body precursors may not survive, or Cas^+^ polar/stalk-like cells may proliferate more. Alternatively or in addition, more cells may adopt a polar/stalk-like fate than a main body fate.

**Figure 4.**
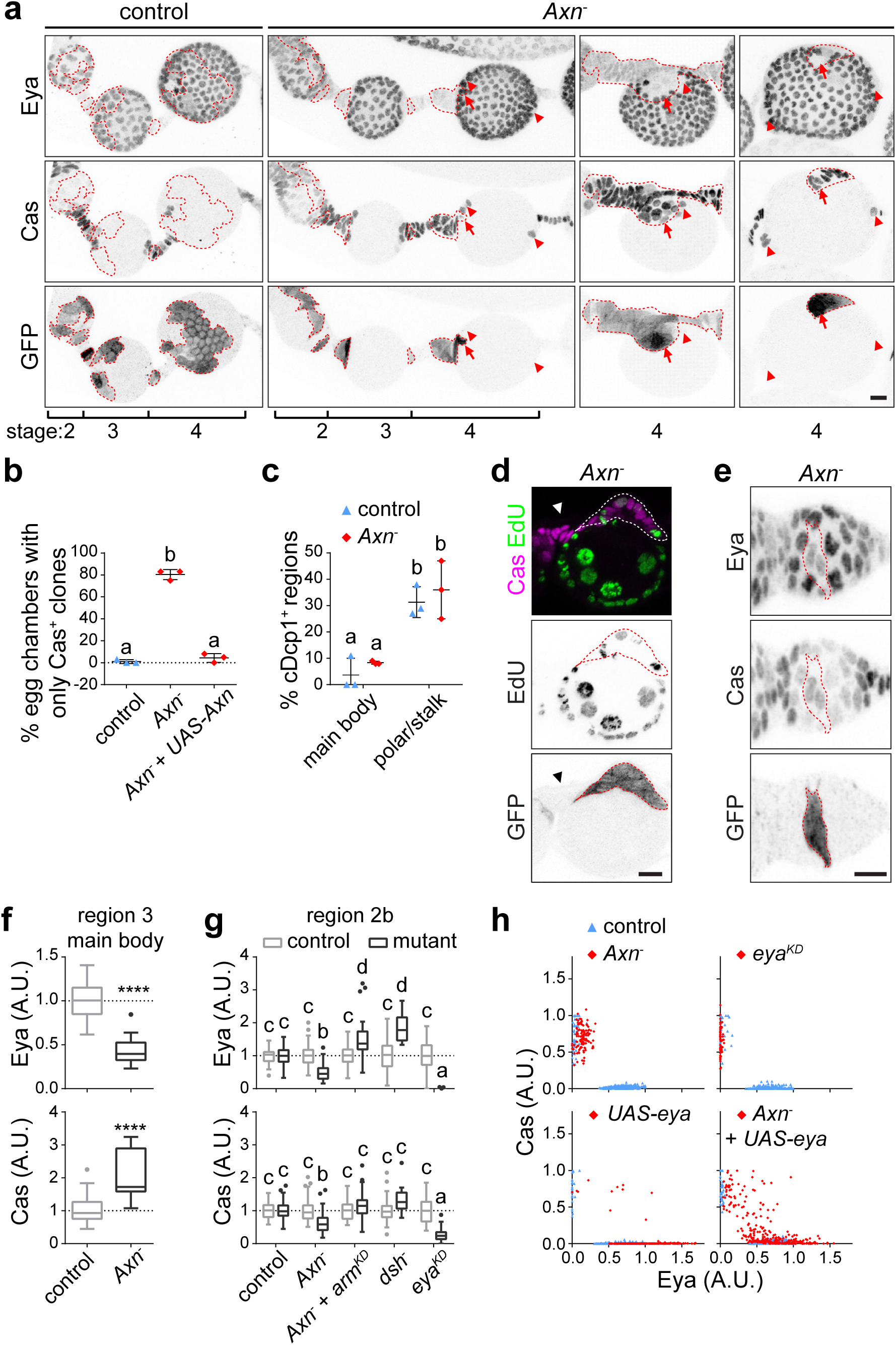
Wnt signalling in differentiating progenitor cells inhibits expression of the main body cell fate determinant Eya. (**a**) 3D projection view of one half of ovarioles with FRT82B control or FRT82B, *Axn^S044230^* mutant (GFP+, dashed lines) FSC clones stained with Eya and Cas. Arrowheads point to control polar cells in stage 4 of *Axn^−^*, and arrows points to the mutant polar cells. Eya^+^ GFP^−^ cells appear in the outlined stage 4 clones due to Z stack projection. (**b**) Quantification of the percentage of stage 3-5 egg chambers with Cas^+^ only clones. Data (mean ± SD) from n = 3 experiments, 77-89 egg chambers per genotype. *p* < 0.0001. (**c**) Quantification of percentage of main body or polar/stalk regions with cells expressing cleaved death caspase-1 (cDcp1^+^) in FRT82B control or FRT82B, *Axn^S044230^* mutant FSC clones. Data (mean ± SD) from n = 3 experiments, 24-36 region 3-stage 2 egg chambers per group. *p* < 0.05. (**d**) Sagittal confocal section of a stage 3 egg chamber with stalk region containing control cells (GFP^−^, arrowhead) compared to *Axn^5044230^* homozygous mutant, ectopic Cas^+^ cells (GFP+, dashed lines). (**e**) 3D projection view of one half of a stage 1 egg chamber with *Axn^5044230^* FSC clones (GFP^+^, dashed lines) stained for Eya and Cas. (**f**) Quantification of Eya and Cas intensity in internal control or *Axn^5044230^* homozygous mutant main body cells in region 3/stage1. Data (median with interquartile range) from n = 3 egg chambers, 59 cells for control, 17 cells for *Axn^−^*. Data were normalized to average Eya or Cas in internal control cells. ****,*p* < 0.0001. (**g**) Quantification of Eya and Cas intensity in germarium region 2b containing FRT82B control, *Axn^5044230^, Axn^5044230^* + *armRNAi, dsh^3^*, or *eyaRNAi* FSC clones. Data (median with interquartile range) from n = 25-100 cells from 3-4 germaria per genotype. *p* < 0.0001. (**h**) Quantification of Eya and Cas intensity in follicle cells with *Axn^5044230^, eyaRNAi, UAS-eya*, or *Axn^S044230^* + *UAR-eya* mosaic FSC clones. Data from n = 1,365-1,434 cells from 4 stage 4 egg chambers per genotype. Scale bar, 10 μm. A.U., arbitrary unit. Samples labelled with different letters are significantly different.

During early stages of oogenesis, apoptosis was common in polar and stalk cells but rare in main body cells (Fig. 4c). *Axn^−^* clones in region 3-stage 2 main body regions did not show a detectable increase in apoptosis (Fig. 4c). The mitotic marker EdU was not detected in control stalk regions after stage 2, but was detected in *Axn^−^* Cas^+^ cells (Fig. 4d), suggesting that these polar/stalk-like cells proliferate more. Strikingly, we found a decrease of Eya and increase of Cas in the main body region in *Axn^−^* cells as early as region 3, suggesting a shift in cell fate (Fig. 4e-f). Such small clones in the main body region likely gave rise to ectopic Eya^−^ Cas^+^ clones observed in stage 4 egg chambers, whereas large main body region clones likely developed into continuous stalks with a single polar cell cluster, causing the egg chamber to appear to bud from the side (Fig. 4a).

To understand how Wnt signalling affects follicle precursor cell differentiation, we quantified the change in Eya and Cas levels in the germarium region 2b. Eya was significantly reduced in the *Axn^−^* clones (Fig. 4g). Conversely, *dsh^−^* cells showed significantly increased Eya in region 2b (Fig. 4g), suggesting that Wnt signalling normally functions to inhibit Eya expression. Reducing Wnt signalling by expressing *armRNAi* in the *Axn*^−^ clone also resulted in increase of Eya, relative to control. Cas was also reduced in *Axn^−^* cells in region 2b, but to a lesser degree than Eya, which was also observed upon *eya* knockdown in this region (Fig. 4g).

Knocking down *eya* in mosaic clones caused all mutant cells to become Eya^−^ Cas^+^, which phenocopied *Axn^−^* (Fig. 4h). To test how important the reduction of Eya is for the fate change in *Axn^−^* cells, we expressed *UAS-eya* in *Axn^−^* clones. Indeed Eya expression restored main body cell fate (Fig. 4h; Supplementary Fig. 7). Therefore Eya is a key target of Wnt signalling. Expression of Eya in *Axn^−^* clones also produced Eya^+^ Cas^+^ cells and Cas^+^ Eya^−^ cells, which were observed when expressing *UAS-eya* alone in mosaic clones. This was likely due to variations in the timing and level of Eya expressions in these experiments, which were not possible to control precisely. These results show that the primary effect of Wnt signalling is to suppress expression of the main body cell fate determinant Eya.

### Notch activity in the germarium promotes polar cell specification

Our results indicate that Wnt signalling enables polar and stalk fates by keeping the Eya level low in the region proximal to the FSC niche, raising the question as to how Wnt relates to Notch in specifying polar cell fate. We used a Notch activity reporter, the *Notch responsive element* (*NRE*) fused green fluorescent proteins ^33^ to address this question. NRE-GFP showed a basal level in all cells, but peaked in a subset of cap cells, in region 2a/2b boundary cells, and in polar cells (Fig. 5a-b). This is consistent with the roles of Notch in the germline stem cell niche ^34,35^, cross-migration of FSC daughters ^8^, and polar cell specification ^9,36^. *Axn^−^* clones showed high Notch activity in polar cells both at normal and ectopic locations that were directly contacting germline, suggesting that a high Wnt signalling predisposes cells to specification as polar cells by Notch (Fig. 5c).

**Figure 5.**
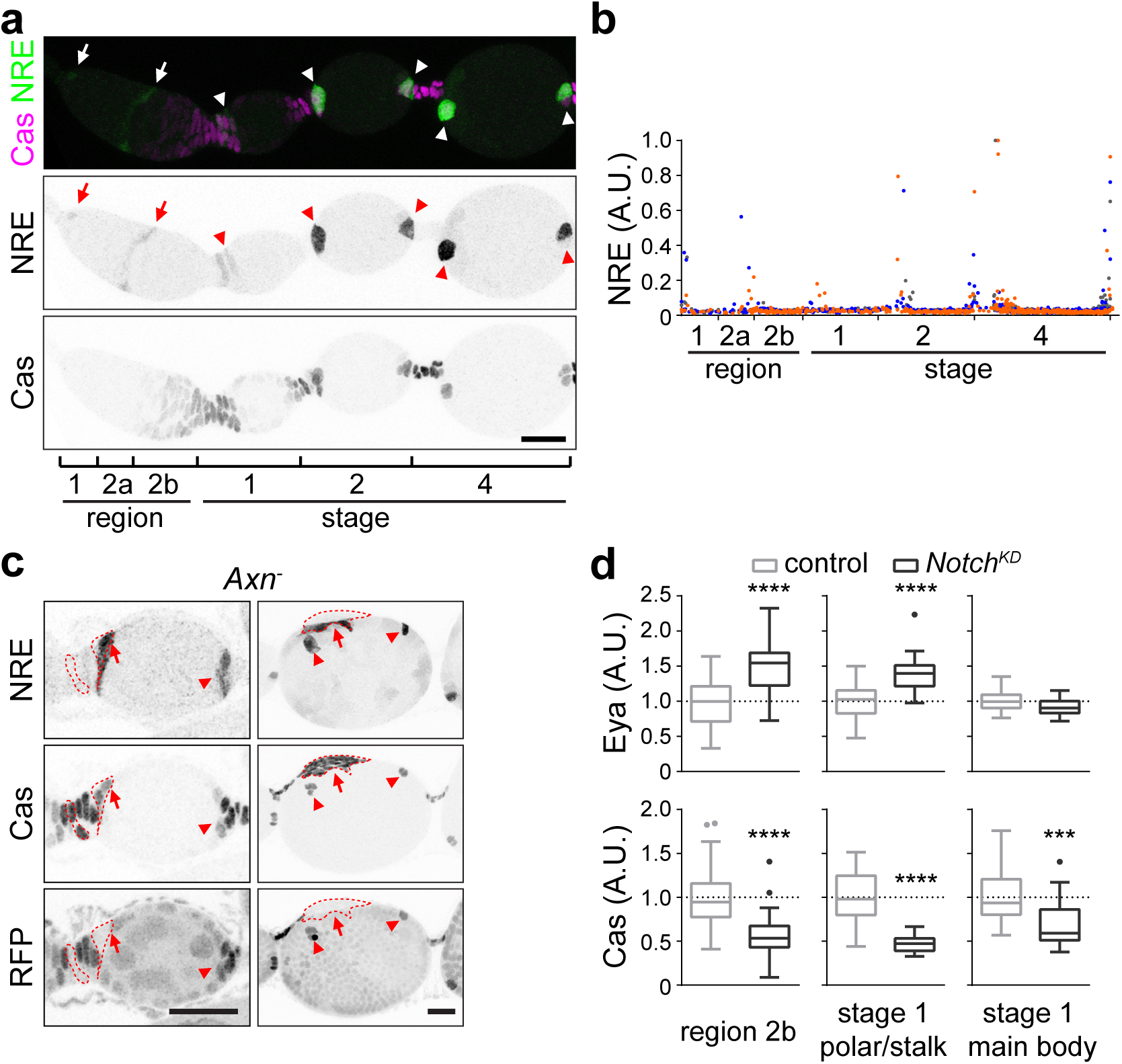
Notch activity in the germarium promotes polar cell specification. (**a**) NRE-GFP pattern from the germarium to stage 4 egg chambers. Arrows point to Notch activity in the cap cells and region 2a/2b boundary cells and arrow-heads point to polar cells. (**b**) Quantification of NRE-GFP intensity in cap cells, escort cells, and follicle cells along the anterior-posterior axis until stage 4. Data from n= 1533 cells in 3 ovarioles. Different colours represent different ovarioles. Data were normalized to maximum NRE-GFP intensity per sample. (**c**) NRE-GFP in *Axn^S044230^* heterozygous control or mutant polar cell clusters in stage 2 (left panel) or stage 6 (right panel) egg chambers. Mutant cells are RFP^−^ (dashed lines). Arrowheads point to control polar cells, and arrows points to the mutant polar cells. (**d**) Quantification of Eya and Cas intensity in germarium containing *notchRNAi* mosaic FSC clones. Data from n = 21-45 cells from 3-4 germaria per region. Data were normalized to average Eya or Cas intensity in internal control cells. ***, *p* < 0.001; ****,*p* < 0.0001. Scale bars, 20 μm. A.U., arbitrary unit.

To assess the effect of Notch signalling on follicle precursor development in the germarium, we quantified the Eya and Cas levels in *NotchRNAi* FSC clones. Reduction of Notch decreased Cas and increased Eya in region 2b as well as the proximal region of stage 1 (Fig. 5d; Supplementary Fig. 8). This is in agreement with the role of Notch in polar cell specification and indeed, we observed high frequency of fused egg chambers when the clone covered the anterior polar cell region. Therefore, Wnt enables polar and stalk fates primarily by suppressing premature Eya expression, whereas Notch promotes polar cell specification and affects both Cas and Eya.

### Hh signalling controls the timing of proliferation versus differentiation

Hyperactive Hh signalling delayed differentiation as shown by the expression pattern of Eya and Cas in stage 4 and later (Fig. 2b,d,f; Supplementary Fig. 9a), as well as expression of the stalk cell marker lamin C (Fig. 6a; ^12^) and the polar cell marker NRE-GFP (Supplementary Fig. 9b), consistent with earlier reports using different markers ^19,30^. How does delayed differentiation change the pattern of follicle cell fates and cause both excess (Fig. 1e; Fig. 6a) and ectopic (Fig. 2d; Supplementary Fig. 9a; ^30,37^) polar/stalk-like cells?

**Figure 6.**
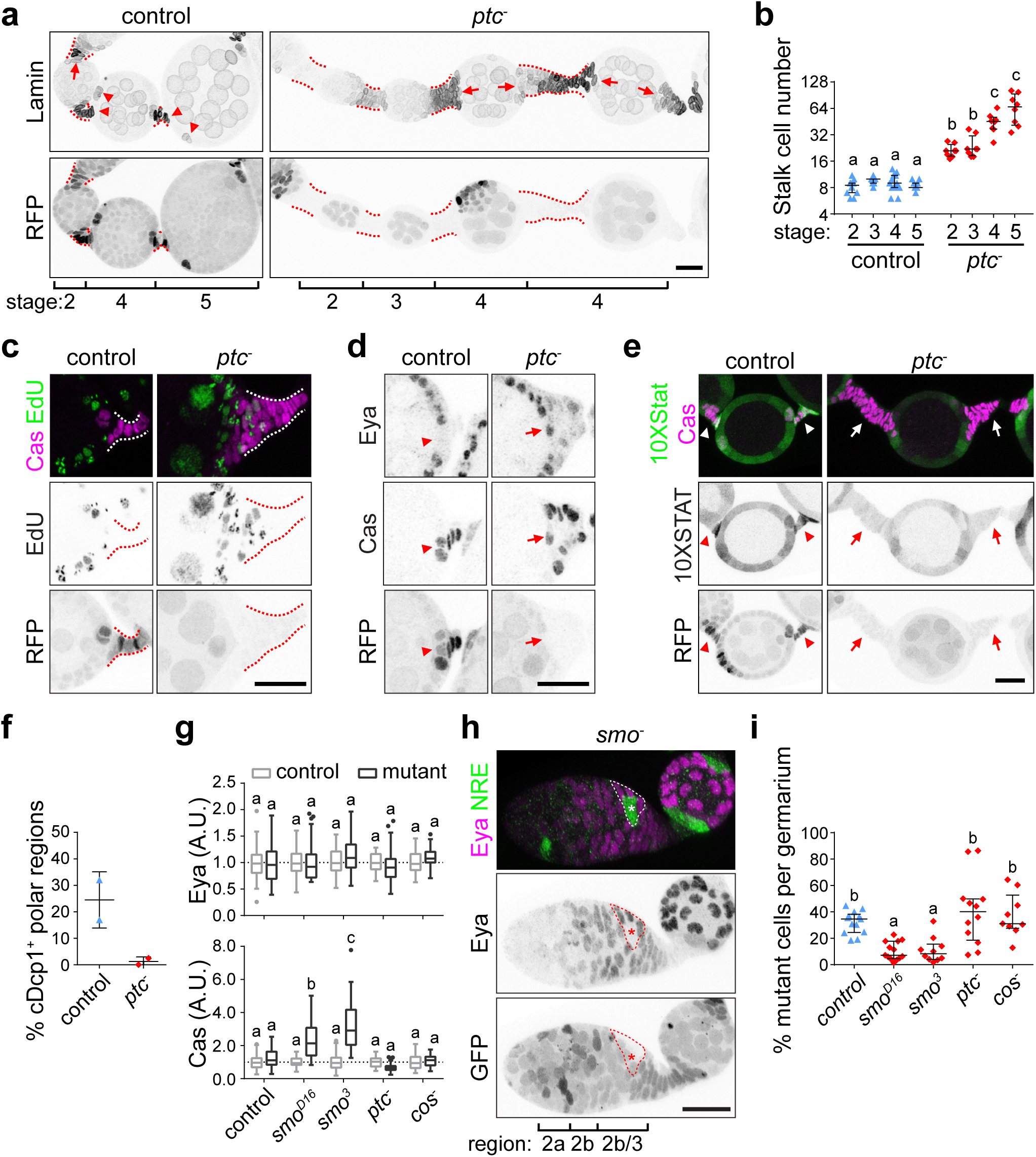
Hh signalling controls the timing of proliferation versus differentiation. (**a**) 3D projection view of ovarioles with *ptc^S2^* heterozygous control or large homozygous mutant (RFP^−^) FSC clones. Dashed lines mark the stalk region. Arrows point to connected polar and stalk regions, and arrowheads point to separated regions. (**b**) Quantification of the number of anterior stalk cells from stage 2-5 egg chambers in *ptc^S2^* heterozygous control or large homozygous mutant FSC clones. Data (median with interquartile range) from n = 8-15 egg chambers. *p* < 0.0001. (**c-d**) Sagittal view of posterior polar/stalk region in *ptc^S2^* heterozygous control or homozygous mutant egg chambers. Dashed lines mark the stalk region. Arrowheads point to control polar/stalk region, and arrows point to the mutant region. (**e**) STAT activity shown by 10XStat-GFP in *ptc^S2^* heterozygous control (arrowheads) or homozygous mutant (arrows) stalk cell regions. (**f**) Quantification of cells with cleaved death caspase-1 (cDcp1+) in *ptc^S2^* heterozygous control or homozygous mutant polar cell regions. Data (mean + SD) from n = 2 experiments, 84-156 stage 2-5 polar cell regions per genotype. (**g**) Quantification of Eya and Cas intensity in germarium region 2b containing FRT40A control, *smo^D16^, smo^3^, ptc^S2^*, or *cos^H29^* mutant FSC clones. Data (median with interquartile range) from n = 21-68 cells from 4-6 germaria per genotype. Data were normalized to average Eya or Cas in internal control cells. *p* < 0.0001. A.U., arbitrary unit. (**h**) NRE-RFP in *smo^3^* mutant FSC clone in a germarium. Mutant cells are RFP^−^ (dashed lines), and the precocious Eya^−^, NRE^+^, presumptive polar cell is marked by *. (**i**) Quantification of mutant clone size in germarium with FRT40A control, *smo^D16^, smo^3^, ptc^S2^*, or *cos^H29^* mutant FSC clones. Data (median with interquartile range) from n = 9-13 germaria,*p* < 0.05. Scale bars, 20 μm. Samples labelled with different letters are significantly different.

We first analysed the long stalk phenotype. The supernumerary stalk cell phenotype appeared most obvious when the majority of follicle cells associated with an egg chamber were mutant. In contrast to control stalks that contained a stable number of stalk cells ranging from 6-13*,ptc^−^* stalks contained an average of 21 cells at stage 2, and continued to increase in number over time (Fig. 6a-b). Another feature of stalk cell maturation is that they normally become physically separated from the polar cells (Fig. 6a, arrowhead), even though stalk cells initially form in the polar region (Fig. 6a, arrow). In contrast, *ptc^−^* stalk cells remained associated with the poles, and with Cas^+^ or lamin C^+^ positive cells on the main body follicle layer (Fig. 6a, arrow). Mature stalk cells with high Cas were normally not proliferative, and thus were EdU-negative after stage 2 (Fig. 6c). In ovarioles with *ptc^−^* clones in regions where stalk was still connected to polar and/or main body cells, Eya and Cas continued to be co-expressed (Fig. 6d), suggesting that they are multipotent precursors. These cells were EdU positive (Fig. 6c). Therefore, we conclude that excessive stalk cell production in *ptc^−^* is due to persistence of precursor cells that contribute cells to the enlarging stalk.

We then asked how supernumerary polar cells form in response to Hh hyper-activation. Typically 3-6 polar cells form and all but two are eliminated by apoptosis ^38^, a process that requires JAK/STAT signalling ^39^. We observed a reduction of JAK/STAT activation in mutant polar cell regions (Fig. 6e), likely caused by the differentiation delay, which also delayed Upd production. There was also reduced apoptosis in the polar cells (Fig. 6f). Therefore, the excess polar cells appeared to result from delayed differentiation, which also postponed apoptosis.

Furthermore, some *ptc^−^* cells in the main body region adopted a polar or stalk cell fate as they matured (Supplementary Fig. 6a), and thus formed ectopic polar/stalk cells. In this situation Eya^+^ Cas^+^ precursors, present due to the developmental delay, differentiated later than normal and thus in the absence of proper spatial patterning signals (see discussion).

If hyperactive Hh signalling delays differentiation, would loss of Hh signalling expedite differentiation? Earlier studies have not detected a defect in differentiation in *smo^−^*^19^, but these studies did not have access to early cell fate markers and quantitative imaging. Using our quantitative analysis of Eya and Cas in the germarium, we observed significantly higher Cas expression in *smo^−^* cells in region 2b (Fig. 6g). We also observed premature differentiation of polar cells using *NRE-RFP* (Fig. 6h), which was never seen in controls at the same stage. The *smo^−^* clone size was significantly smaller than *ptc^−^* or control, suggesting premature termination of proliferation (Fig. 6i).

### Wnt and Hh double mutants show additive effects on egg chamber patterning

Since Wnt and Hh show independent activities and have distinct functions in follicle precursor cell differentiation, we hypothesized that combined loss of Wnt and Hh activity should show a more extreme egg chamber formation phenotype than either alone. We used C306-Gal4 to drive *smo* and *dsh* RNAi in the follicle precursor cells (Supplementary Fig. 10a). Indeed, we observed an increase in the frequency of egg chamber fusions to 80% in double RNAi knockdowns compared to 30-40% for the single knockdowns (Fig. 7a; Supplementary Fig. 10b), a defect indicating a problem with producing the correct number of polar and stalk cells in the right location for egg chamber budding.

**Figure 7.**
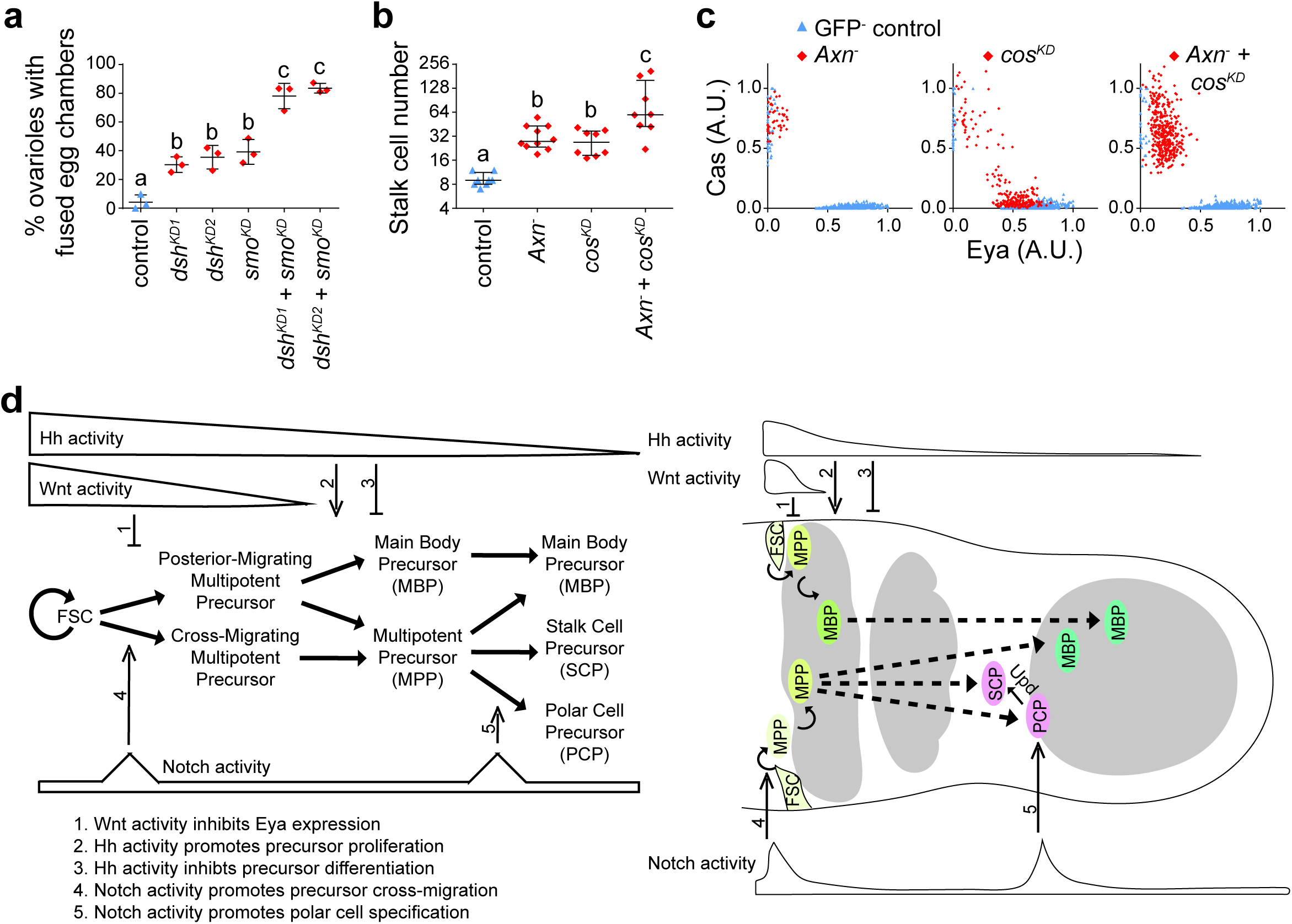
Wnt and Hh exert independent effects in egg chamber patterning. (**a**) *C306-Gal4* driven knockdown of *dsh* and/or *smo* in follicle precursor cells. Data (mean ± SD) from n = 3 experiments, 145-189 ovarioles per genotype. (**b**) Quantification of stalk cells anterior to stage 4-5 egg chambers in FRT82B control, *Axn^S044230^, cosRNAi*, or *Axn^S044230^* + *cosRNAi* mosaic FSC clones. Data (median with interquartile range) from n = 7-9 egg chambers per genotype. (**c**) Quantification of Eya and Cas intensity in follicle cells with *Axn^S044230^, cosRNAi*, or *Axn^S044230^* + *cosRNAi* mosaic FSC clones. Data from n = 890-1,017 cells from 3 stage 4 egg chambers per genotype. A.U., arbitrary unit. (**d**) Model for Wnt, Hh, and Notch in follicle precursor cell differentiation in the germarium. Samples labelled with different letters are significantly different at *p* < 0.01.

We further predicted that hyper-activation of both Wnt and Hh would show an additive effect. Indeed, whereas there are normally 8-10 stalk cells, we observed an average of 30 stalk cells in *Axn* or *cos* single mutant, and a number that doubled following Wnt and Hh double hyper-activation (Fig. 7b; Supplementary Fig. 11). The *Axn* or *cos* double mutant cells are still largely biased toward Cas^+^ polar/stalk like cells (Fig. 7c), suggesting that the Eya^+^ Cas^+^ precursors that accumulated due to delayed differentiation preferentially adopted a polar/stalk fate in the presence of high Wnt signalling.

## Discussion

When adult tissue stem cells divide asymmetrically to self-renew and produce a daughter cell that commonly becomes a transit amplifying precursor, one possibility is that the stem cell retains its character by virtue of its association and proximity to niche signals whereas the transient amplifying precursor acquires its properties due to displacement from the niche. Here we use the Drosophila ovary model to address the roles of secreted niche factors in diversification of precursor cell fates as they leave the niche. We found that Wnt and Hh act as morphogens that not only maintain FSC at a high concentration, but also provide critical instructions for the development of the transit amplifying follicle precursor population at a lower concentration. The distinct shapes of the gradients of these niche signals serve important functions. A key finding is that Wnt signals maintain multipotency in a subset of precursors that maintain closer proximity to the niche, whereas reduced Wnt signalling activates Eya expression in precursors displaced further from the niche and this restricts them to the main body cell fate.

Earlier studies reported egg chamber formation defects due to reducing the Wnt or Hh ligand levels ^12,18,30^. However, recent studies found roles of Wnt and Hh in escort cells to affect germline differentiation ^16,17,40,41^, therefore the ovariole defects from reduction of Wnt or Hh ligands could either be caused by effects on the germline and/or follicle cells. We clarified this issue by reducing Wnt and Hh intracellular signalling components directly in follicle cells in mosaic clones, or by RNAi knockdown specifically in follicle precursor cells.

In the germarium, the follicle precursor cells in region 2b and region 3 contain both specified and unspecified cells ^8^, yet previous studies lacked cell fate markers and quantitative methods to assess the influence of different signalling inputs. Our quantitative analysis of the Wnt, Hh and Notch signalling reporters as well as the differentiation factors, Eya and Cas, not only provide a sensitive and detailed description of the FSC differentiation process, but also reveal early changes in the germarium that were not previously detected. First, reduction of Wnt activity by *dsh^−^* or *armRNAi* increased the expression of Eya, a key main body fate determinant ^21^, and reduction of Wnt by *C306-Gal4* driven *dshRNAi* in large numbers of precursor cells caused egg chamber fusing, a phenotype also seen when driving *UAS-eya* by *C306-Gal4* ^25^. Second, reduction of Hh activity by *smo^−^* resulted in increased levels of Cas in region 2b, indicative of premature differentiation. However, the Eya level did not increase in region 2b, which is likely because of inhibition of Eya by Wnt in this region. Third, reduction of Notch activity shifted the balance between Eya and Cas in region 2b.

Our results suggest an integrated model for how Wnt and Hh stem cell niche signals act in coordination with the differentiation factor Notch to establish the initial asymmetry and specify cell fates during follicle precursor cell differentiation in the germarium (Fig. 7d). We propose that follicle precursor cells expressing low levels of Cas and Eya have intrinsically unstable fates that can stochastically give rise to Eya^+^ Cas^+^, Eya^+^ Cas^−^, or Eya^−^ Cas^+^ cells in the absence of specific instructive signals. The spatial patterns of Wnt, Hh and Notch function as environmental cues to direct these precursor cells to produce the appropriate numbers and types of differentiated cells.

We show that hyper-activation of Wnt signalling biases follicle cells to polar and stalk cell fates, both of which lack Eya and express high levels of Cas. A simple interpretation of these results might suggest that the follicle precursor cells that retain higher Wnt signalling adopt a polar/stalk precursor fate whereas the ones further displaced from the niche with disinhibited Eya adopt the main body precursor fate. However there are clear data demonstrating that there is no dedicated polar/stalk precursor ^8^, while the same clonal analyses are perfectly consistent with the existence of a dedicated main body precursor cell. Therefore we propose that the steep gradient of Wnt in region 2b results in progenitor cells with different levels of Wnt signalling. The progenitor cells that are displaced from the niche, the posterior daughter of the FSC, escape Wnt signalling and therefore express Eya and adopt a dedicated main body precursor fate. The progenitors that remain closer to the niche, the cross migrating cell and its daughter, receive more Wnt ligand ^16,17^ and thus maintain a pluripotent state that is permissive for polar and stalk fates but not irreversibly committed to those fates. Both the absolute and relative levels of Eya and Cas are likely to be important for the final fate.

The Hh gradient in the germarium provides important temporal information to be coupled with the spatial information as developing cysts move posteriorly and away from the FSC niche. In the absence of proper coordination of differentiation and fate, cells can acquire random fates, as observed in stage 4 *ptc* or *cos* mutant clones for example (Fig. 2d, f). These cells lack the spatial patterning normally provided by Wnt signalling by the time they differentiate. When Wnt and Hh signalling are co-activated, most of the cells adopt polar/stalk-like fate (Fig. 7c).

Notch specifies polar cells through two actions in the germarium. First, Notch activity affects expression of Eya and Cas in region 2b, likely indirectly by promoting the cross migration of FSC daughters ^8^ so the precursor cells remain in high Wnt for longer. As a result, cells with reduced Notch activity in region 2b preferentially undergo posterior rather than lateral migration, and therefore escape from Wnt sooner and express higher Eya. Meanwhile, the reduction of Cas in Notch mutant cells is likely because they differentiate slower and therefore escape from Hh later, similar to the Notch and Hh antagonism described in the mitosis to endocycle switch ^42^. Second, relatively higher Notch activity due to germline contact ^43^ and fringe expression ^44^ specify polar cells from the Eya^low^ multipotent precursor cell pool. Although essential for polar cell specification, hyper-activation of Notch does not seem sufficient for polar cell formation since excess polar cells only form in the two poles rather than on the main body region ^10^, This implies that additional spatial information is required besides Notch activity for polar cell formation. Here we found that spatial information to be a short-range Wnt signalling. Notch activity coordinates with Wnt to specify polar cells.

Wnt, Hh and Notch are common players in many adult stem cell systems including the skin, gut, and blood ^45–47^, which all possess a transit-amplifying progenitor pool close to the stem cell niche. Our finding of the distinct functions of Wnt, Hh, and Notch in precursor cell fate specification provides an integrated model for how multiple signalling inputs establish the initial asymmetry in cell fates. The additive effect of hyper-activation of multiple signalling pathways that we observed may have implications in cancer stem cell and its targeted treatment ^48^.

## Methods

### Drosophila genetics and mosaic clone induction

Fly strains used in this study are listed in Supplementary Table 1. Fly genotypes used in each experiment are listed in Supplementary Table 2. Stocks were maintained at room temperature. Crosses were initiated at room temperature and transferred to 25 °C at 2-3 instar larvae stage. For C306Gal4; tubGal80ts experiments, adult female flies were transferred to 29 °C for 7-10 days after eclosion. Egg chamber stage was determined based on germ cell nucleus diameter listed in Supplementary Table 3.

**Supplementary Table 1:**
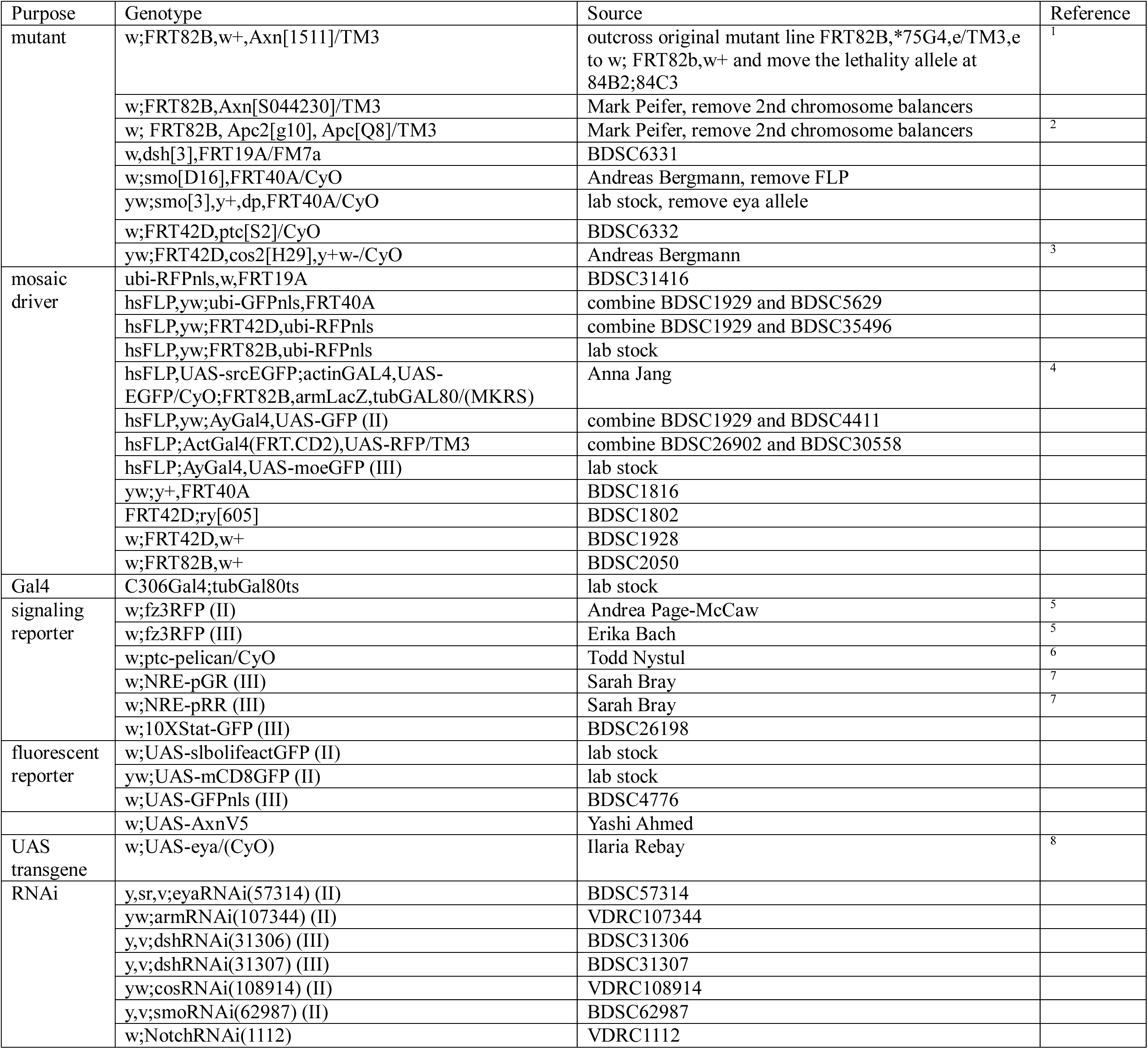
List of fly stains used in this study

**Supplementary Table 2:**
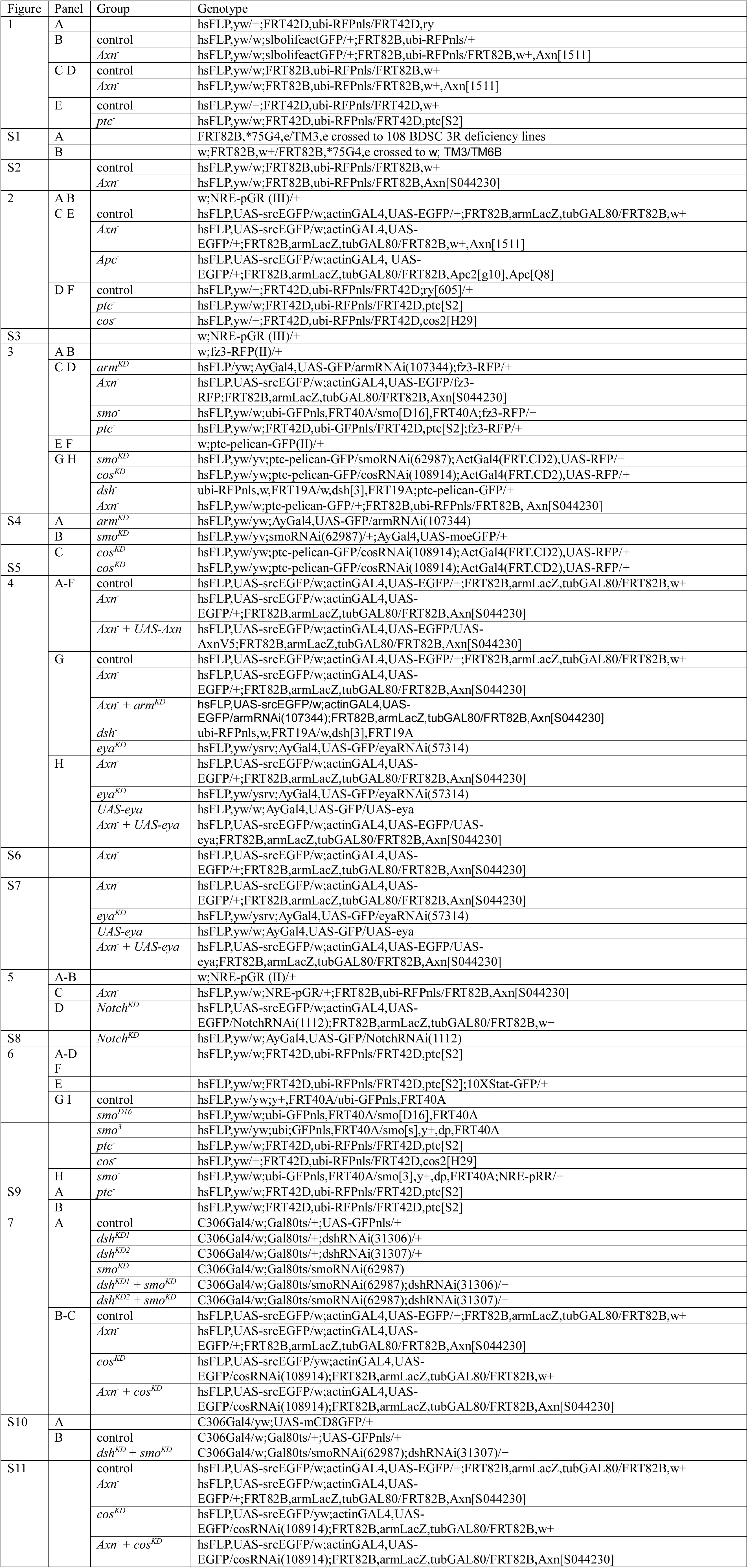
List of fly genotypes used in each experiment

**Supplementary Table 3:** Stage of egg chamber development

| Stage* | Nurse cell nucleus diameter (µm) |  |
| --- | --- | --- |
|  | anterior | posterior |
| 2 | 5-7 |  |
| 3 | 7-9 |  |
| 4 | 9-12.5 |  |
| 5 | 12.5-15.8 |  |
| 6 | 15.8-18 |  |
| 7 | 18-21.8 | 23-30 |
| 8 | 21.8-25 | 28-32 |
| 9 | 25-35 | 32-43 |
| 10 | 35-45 | 42-50 |

\* Stage according to <sup>9</sup>

Mosaic clones were generated using the FLP/FRT system. 8-9 newly eclosed adult female flies (1-2 days old) along with 8 males were collected in a vial with wet yeast paste (dry yeast and water 1:1.5) and dry yeast and kept at 25 °C. Flies were flipped without CO_2_ to a fresh vial daily until dissection, and heat shocked 2 days after collection. Males were added if less than 3 were present to ensure optimal ovary development. For making FSC clones up to stage 5, flies were heat shocked twice for 1 hour, about 4 hours apart, in a 37 °C water bath, and then were kept at 25 °C for 5-7 days before dissection. For RNAi knockdown experiments, flies were transferred to 29 °C after heat shock (except for *eyaRNAi* and *armRNAi*, which were kept at 25 °C). For making FSC clones up to stage 8, flies were kept for 6-8 days before dissection. For border cell clones in Fig. 1d, flies were heat shocked once for 30 minutes and kept for 4-5 days before dissection. For MARCM clones, we observed some leaky GFP expression in follicle cells in stage 6 and later, likely due to actinGal4 being too strong in stage 6 and later such that tubGal80 was not able to suppress all Gal4 activities. Therefore, we only analysed MARCM clones before stage 6.

### Immunostaining and EdU incorporation

Adult female ovaries were dissected in Schneider’s *Drosophila* medium (Thermo Fisher Scientific, Waltham, MA) with 20% fetal bovine serum and transferred to a 0.6 ml microfuge tube with 100 μl dissection medium. Ovaries were dissociated by pipetting up and down approximately 50 times using a 200 μl pipette set to 50 μl Dissociation in this way causes random physical damage to the egg chambers ^49^, but we found it more efficient than pulling ovarioles out of the muscle sheath using forceps, which causes more damage to the germarium or younger egg chambers. Ovarioles were immediately fixed for 20 minutes in 4% paraformaldehyde at 4 °C. After fixation, ovarioles were washed with PBS/0.4% Triton X-100 (PBST), and then incubated with primary antibodies overnight at 4 °C. The following day, ovarioles were washed with PBST before incubation in secondary antibody for 1.5-2 hours. After removal of secondary antibodies, samples were stained with Hoechst for 20 minutes. Samples were washed in PBST before sorting in PBST. Sorting was conducted by using forceps under a dissection microscope to remove mature eggs and clustered ovarioles from a given sample for optimal mounting. Without sorting, mature eggs make it difficult to compress the sample, the germaria can be tilted, whereas clustered ovarioles often overlap each other rendering imaging difficult. After sorting, samples were stored in VECTASHIELD (Vector Laboratories, Burlingame, CA) at 4 °C.

The following antibodies were used in this study: chicken anti-GFP (1:2000, Abcam, Cambridge, UK; 13970) (used to amplify MARCM GFP, Flipout GFP, Ptc-pelican-GFP, and NRE-GFP, not used on negative mosaic ubi-GFPnls), rabbit anti-dsRed (1:1000, Takara Bio USA, Mountain View, CA; 632496) (used to amplify Flipout RFP and NRE-RFP, not used on negative mosaic ubi-RFPnls or Fz3-RFP), mouse anti-Eyes Absent [1:50-200, Developmental Studies Hybridoma Bank (DSHB), Iowa City, IA; 10H6, needs pre-absorption if staining is noisy], mouse anti-Fascillin III (1:50, DSHB 7G10), rat anti-E-cadherin (1:50, DSHB DCAD2), rabbit anti-Castor [1:5000, Ward F. Odenwald, ^50^], mouse anti-Armadillo (1:100, DSHB N27A1), mouse anti-Smoothened (1:4, DSHB 20C6), rat anti-Cubitus interruptus (1:10, DSHB 2A1), rabbit anti-cleaved Drosophila caspase 1 (1:200, Cell Signaling Technology, Danvers, MA; 9578), mouse anti-Notch intracellular domain (1:200, DSHB C17.9C6), and mouse anti-Lamin C (1:200, DSHB LC28.26).

For 5-ethynyl-2’-deoxyuridine (EdU) incorporation, adult female ovaries were dissected in Schneider’s *Drosophila* medium with 20% fetal bovine serum and transferred to a microfuge tube with the dissection medium plus 40 μM EdU, and kept at room temperature on a shaker for 1 hour. Ovarioles were then dissociated, fixed, and stained with primary and secondary antibodies as described above. Before staining with Hoechst, an EdU detection reaction was performed according to the manufacturer’s manual (Thermo Fisher Scientific).

### Imaging and image processing

Due to the spherical organization of the egg chambers, few follicle cells have their nuclei located on the same imaging focal plane. Therefore, we imaged the egg chambers in full Z stacks. Samples were mounted on a glass slide in VECTASHIELD (25 μl for early stage ovarioles, or 65 μl for stage 9/10) using a 22 mm X 40 mm cover glass, to ensure that the germarium was mounted flat, but not compressed, and that later stages were compressed to a consistent degree. All images were taken on a Zeiss LSM780 confocal microscope, using a 40x 1.4 N.A. oil objective. Z stacks covering the entire germaria or ovarioles were taken with a 0.43 μm step size for germarium and ovarioles, or a 1 μm step size for border cell clusters. XY resolution is 0.14 μm for germaria, or 0.35 μm for ovarioles. Laser power corrections were applied by increasing the laser power as the objective scans from the top of the sample to the bottom of the sample, so that the signal on the bottom did not appear weaker than the top. 3D images were visualized in Imaris (Bitplane, South Windsor, CT), and annotated in Excel (Microsoft, Redmond, WA), to categorize the developmental stage, sample condition, mounting condition, imaging condition, clone location, and result interpretation. Developmental stage was determined as described above. Sample condition includes whether they were damaged, or still tightly packed in muscle sheath. Severely damaged egg chambers had an incomplete follicle epithelium and leaky germ cells, or large patches of follicle cells without nuclear stain. Mild damage caused a small patch of follicle cells to show condensed Hoechst staining, and diffused or reduced nuclei Eya, Cas, or ubi-GFP/RFPnls signal ^49^. Samples with severe damage were not analysed, and the damaged cells in a sample with mild damage were not included in the analysis. We preferred to analyse samples out of the muscle sheath, because their morphology was not affected by squeezing from neighbouring egg chambers. Samples tightly packed in muscle sheath were not used for intensity measurement because it was difficult to perform laser power correction. Mounting condition denotes if the sample was too compressed or too tilted. If the germarium was too compressed, the germline cysts were squeezed and it was difficult to perform 3D rotation as described below. If too tilted, laser correction became difficult. Imaging condition marks whether the image was taken with proper laser power correction. This was estimated by comparing the signal intensity of the top, middle, and bottom of the sample visually, and was quantified as described below. Clone location and result interpretation were listed to help summarize the results, draw conclusions based on the phenotype seen across multiple ovarioles, and select representative images for presentation.

Representative images were exported from Imaris using either Easy 3D view or slice view. Since different follicle cell nuclei were located on different focal planes, 2-5 μm Z stacks were used to show single follicle cell layers, while 12-25 μm Z stacks were used to show one half of the egg chambers. Exported images were rotated and cropped in Photoshop (Adobe, San Jose, CA). Single channel images were converted from a black background to a white background using Invert LUT function in Fiji ^51^.

### 3D quantification

Image segmentation was performed using Imaris. First, samples were rotated using the Free Rotate function. Egg chambers were rotated so the polar cells aligned horizontally, with the anterior to the left. Germaria were rotated in two steps. The first step positioned region 2b cysts vertically in the Z-direction by placing an Oblique Slicer in the mid-sagittal section of the germaria, and performing free rotation to the orthogonal view of the oblique slicer. The second step rotated the germaria anterior to the left to place region 2b cysts vertically in XY-direction. Second, follicle cell nuclei were detected using the Spots function. For the germaria, a 2.5 μm diameter spot size was used for automatic spot detection in the channel with follicle cell nuclei signals. Spots were then manually edited so that each follicle cell was marked. Dividing, dying, or damaged cells showed clear signs, including condensed Hoechst staining and diffused, or reduced, nuclei Eya, Cas, or ubi-GFP/RFP signal, and were not quantified. The 2.5 μm spots were then used to create a masked channel, and automatic spots detection based on that channel was applied to create 1.75 μm spots, so that only the center of the nuclei with a strong and even signal was used for quantification. For egg chambers, a 3.46 μm diameter spot size was used for automatic spot detection, followed by reduction to 1.75-2 μm. Third, background intensities were estimated by placing 8-12 1.75-2 μm spots in two Z planes in same region as the measured follicle cells. For Eya and Cas, background spots were placed in the germ cell cytoplasm, while for Wnt or Hh reporters they were placed in the region 2b germ cell nuclei. Fourth, accuracy of laser power correction was determined by selecting control cells at the top, middle, and bottom of the germarium or egg chamber in the same region, and comparing their signal intensities.

Data for spot position and channel mean intensity were exported from Imaris, and processed using MATLAB (MathWorks, Natick, MA) for background subtraction, comparison of top, middle, and bottom intensity, and normalization, and plotted using Prism (GraphPad, La Jolla, CA).

### Statistics

Statistics were performed using Prism. Unpaired t-test was used for comparing two groups, and Ordinary one-way ANOVA, followed by Tukey’s multiple comparisons test, was used for comparing multiple groups. For box plots, the Tukey method was used for plotting whiskers and outliers.

## Acknowledgements

This work was supported by NIH grant GM46425 to D.J.M. We thank Abby Padgett, Haley Burrous, Benjamin Cheng, Sreesankar Easwaran, and Junjie Luo for technical assistance. We thank Dr, Ward F. Odenwald and the Developmental Studies Hybridoma Bank for providing antibodies and Drs. Yashi Ahmed, Erika Bach, Andreas Bergmann, Sarah Bray, Anna Jiang, Todd Nystul, Andrea Page-McCaw, Mark Peifer, Ilaria Rebay, the Bloomington Drosophila Stock Center, and the Vienna Drosophila Resource Center for providing fly stocks. We acknowledge the use of the NRI-MCDB Microscopy Facility and the Imaris computer workstation supported by the Office of The Director, National Institutes of Health of the NIH under Award # S10OD010610.

## Author Contributions

W.D., A.P. and T.K. performed the experiments. W.D., A.P. and D.J. M. prepared the manuscript. All authors participated in the design of the experiments, the interpretation of the data, and the production of the final manuscript.

## Competing Financial Interests statement

The authors declare no competing final interests.

---

**Supplementary Figure 1.**
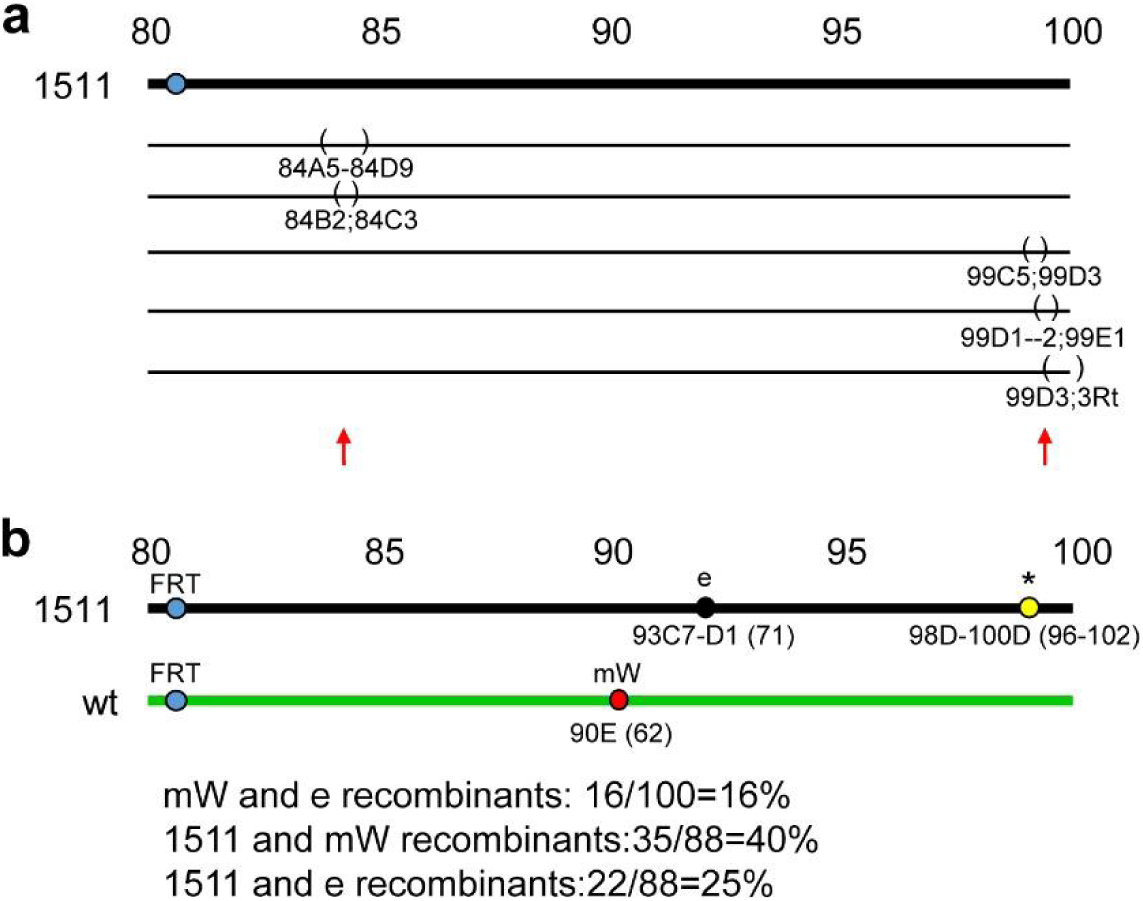
Genetic mapping of *Axn^1511^*. (**a**) Mapping of *Axn^1511^* by crossing to the chromosome 3R deficiency kit. Lethality was found when crossing 1511 to the 5 deficiency lines with the indicated deletions (brackets), which suggests that 1511 contains two lethal mutations (arrows). (**b**) Recombination mapping of *Axn^1511^* by crossing to FRT82B, mW90E. The large border cell cluster phenotype (1511), red eye color (mW), and dark body color (e) were scored in recombinant lines. 1511 was mapped to chromosomal position 96-102 as estimated by recombination rates. The lethal mutation at chromosomal position 84 was removed in the recombinant used in Figure 1 and 2.

**Supplementary Figure 2.**
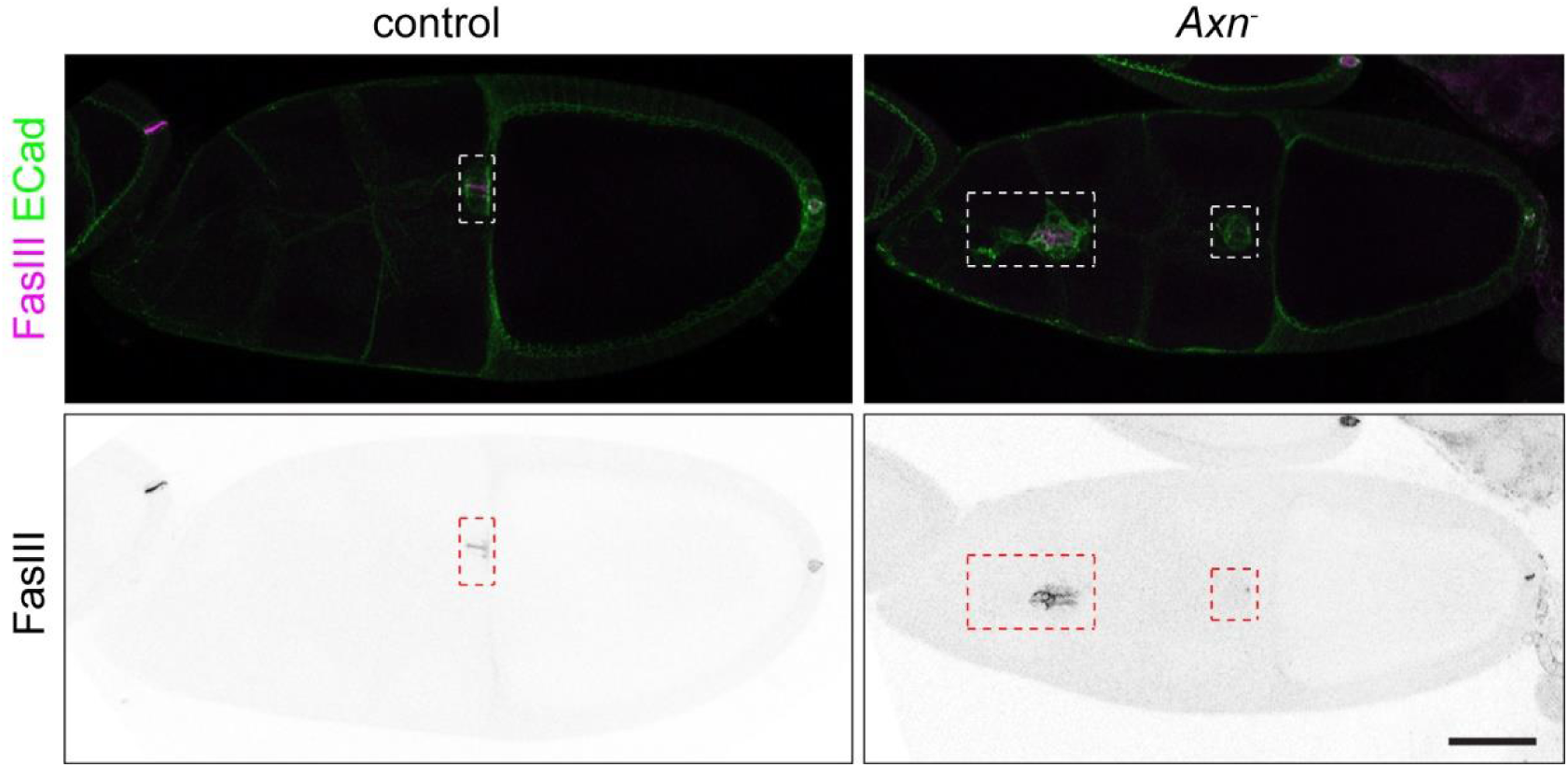
*Axn^S044230^* mutant clones cause supernumerary polar cells. Border cell clusters (dashed boxes) in FRT82B control or FRT82B, *Axn^3044230^* mosaic stage 10 egg chambers. ECad enriches in border cell clusters and FasIII accumulates on polar cell membrane. Scale bar, 50 μm.

**Supplementary Figure 3.**
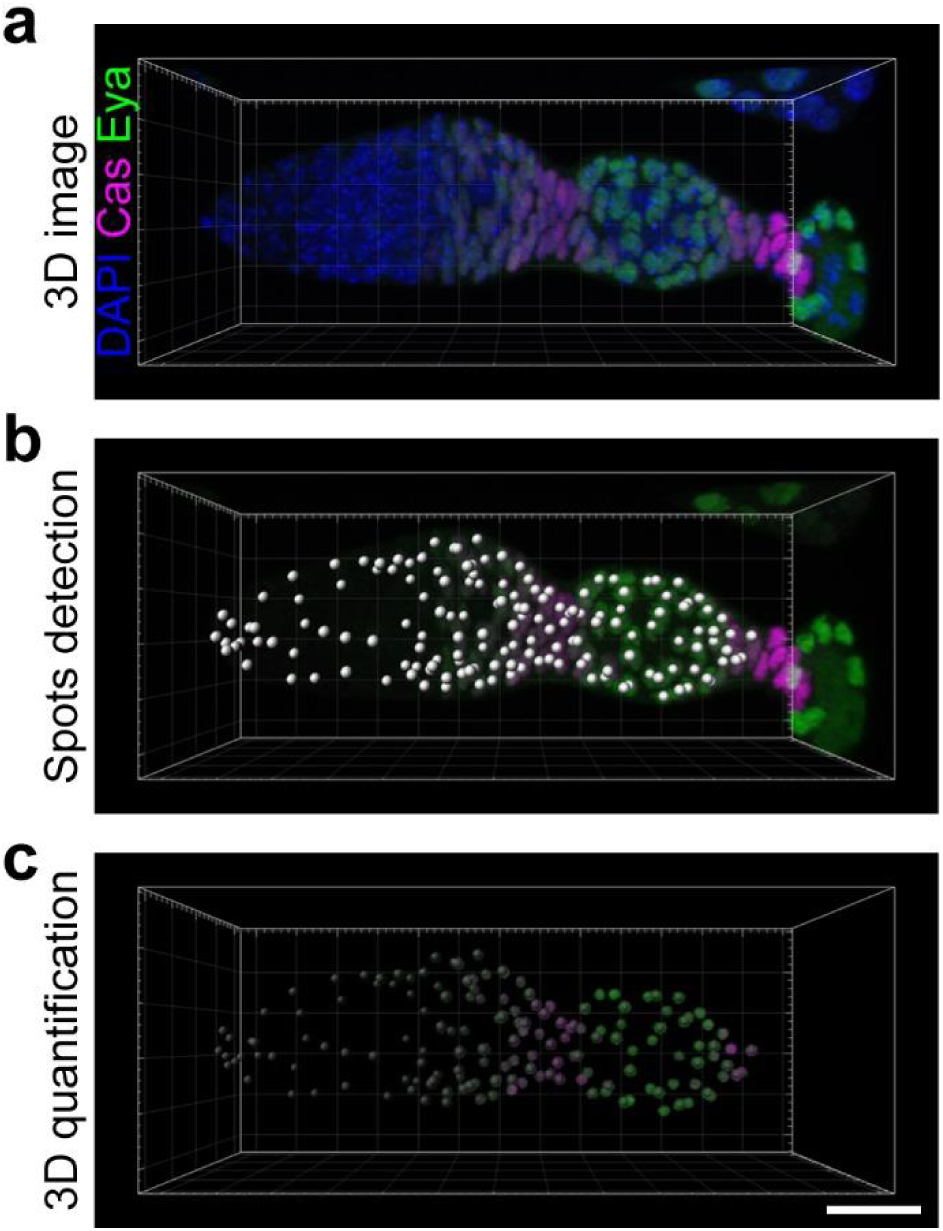
Example of 3D quantification of the levels of both Eya and Cas in every somatic cell in a germarium. (**a**) Image in 3D view. (**b**) Nuclei of somatic cells were first automatically detected by 2.5 μm spots (white dots) using the Eya channel, followed by manual proof editing using the Eya and DAPI channels to ensure one spot per nucleus. (**c**) A masked channel was created by setting the outside of the 2.5 μm spots to 0 intensity, and a 1.75 μm spot (semi-transparent dots) was automatically placed in the center of the 2.5 μm spot to get a strong and even nuclei intensity measurement. See methods for further details. Scale bar, 20 μm.

**Supplementary Figure 4.**
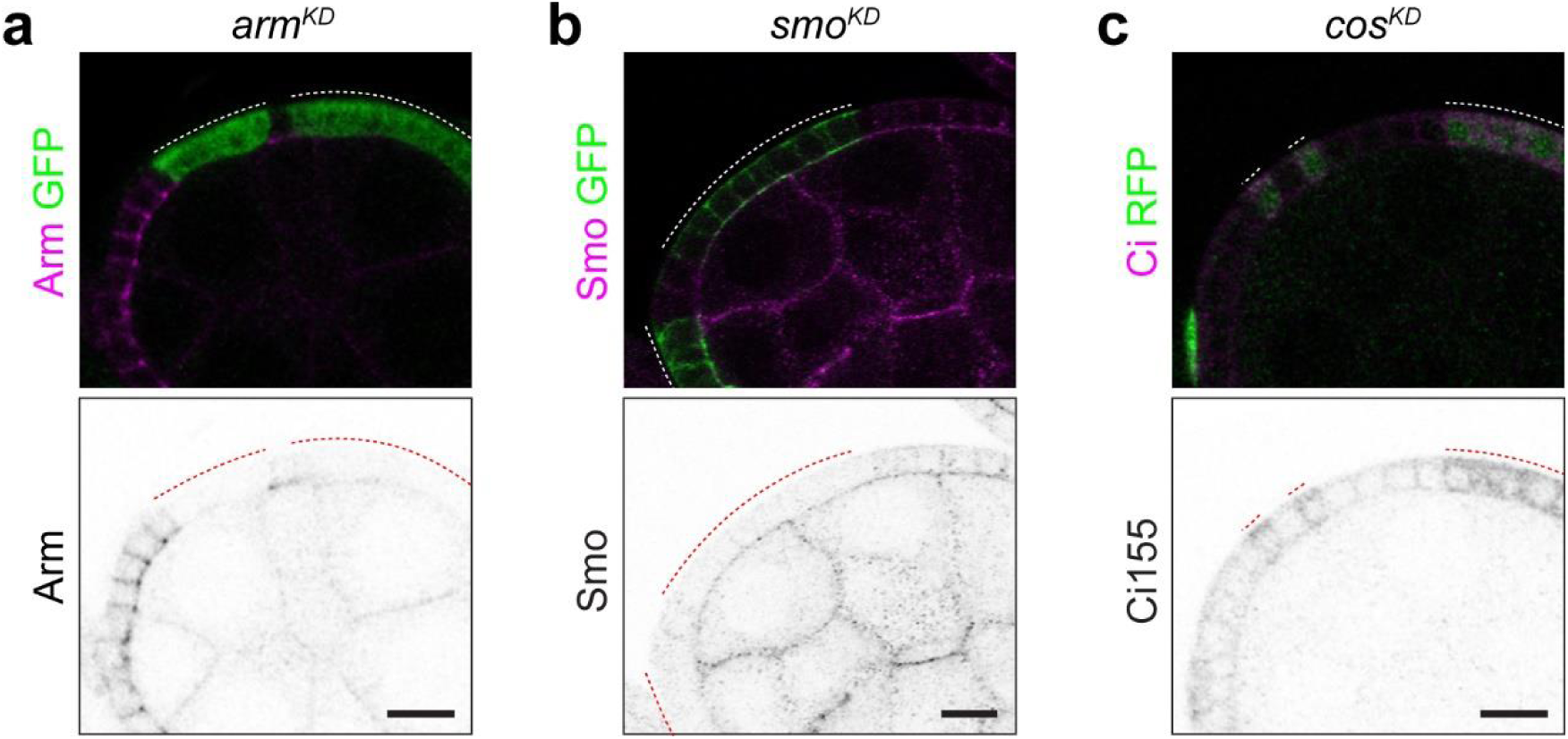
Supplementary Figure 4. Validation of RNAi lines used. (**a**) Sagittal confocal section of *armRNAi* flip-out clones (GFP^+^, indicated by the dashed line) in a stage 5 egg chamber stained with anti-Arm antibody (magenta in top panel, black in bottom). (**b**) Sagittal confocal section of *smoRNAi* flip-out clones (GFP^+^) in a stage 6 egg chamber stained with anti-Smo antibody (magenta in top panel, black in bottom). (**c**) Sagittal confocal section of *cosRNAi* flip-out clones (RFP^+^) in a stage 5 egg chamber stained with anti-Ci155 antibody (magenta in top panel, black in bottom). Scale bars, 10 μm.

**Supplementary Figure 5.**
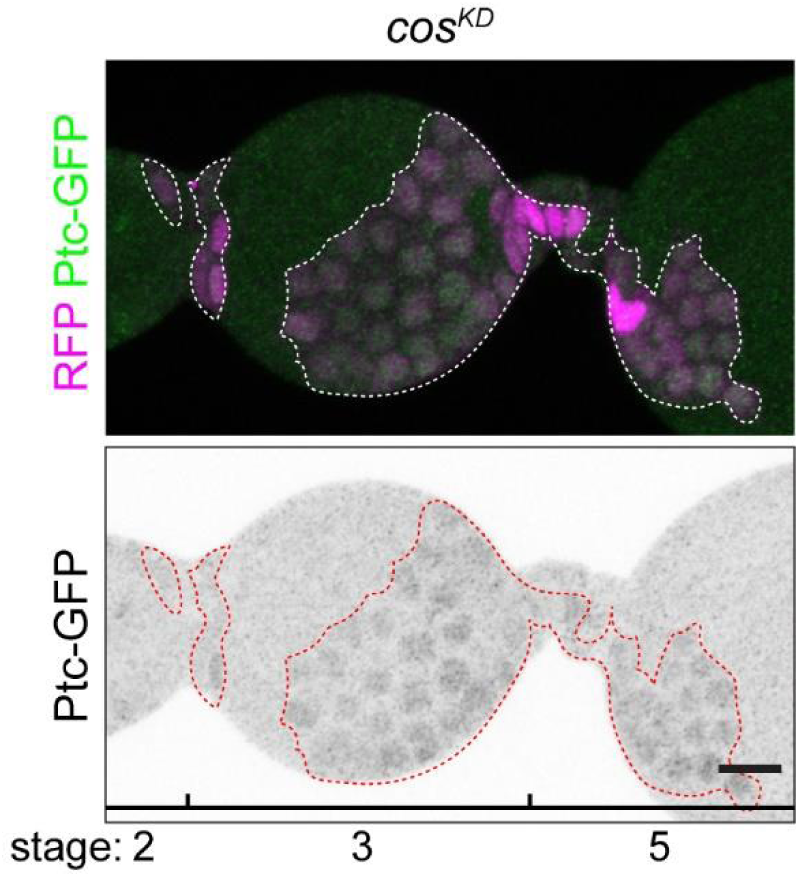
*ptc-GFP* signal in *cos^KD^* ovariole. 3D projection view of one half of stage 3-5 egg chambers with *cosRNAi* flip-out clones (RFP+) showing the effect on expression of the *ptc-GFP* reporter. Scale bar, 10 μm.

**Supplementary Figure 6.**
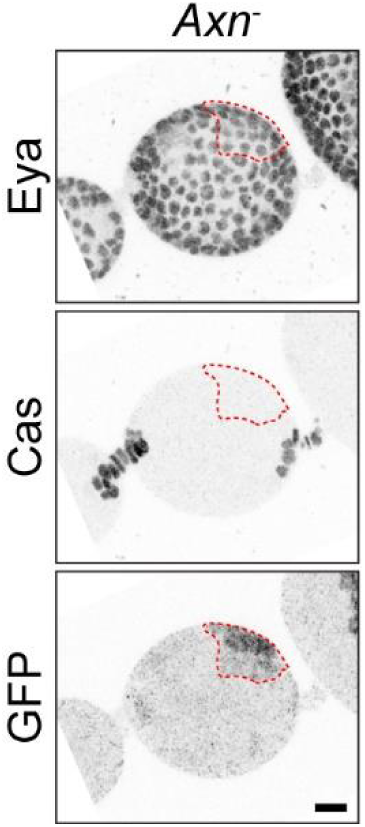
*Axn^S044230^* transient clone does not bias towards polar/stalk fate. 3D projection view of one half of egg chambers with *Axn^S044230^* transient clones 2 days post clone induction (GFP+, dashed lines) stained for Eya and Cas. Scale bar, 10 μm.

**Supplementary Figure 7.**
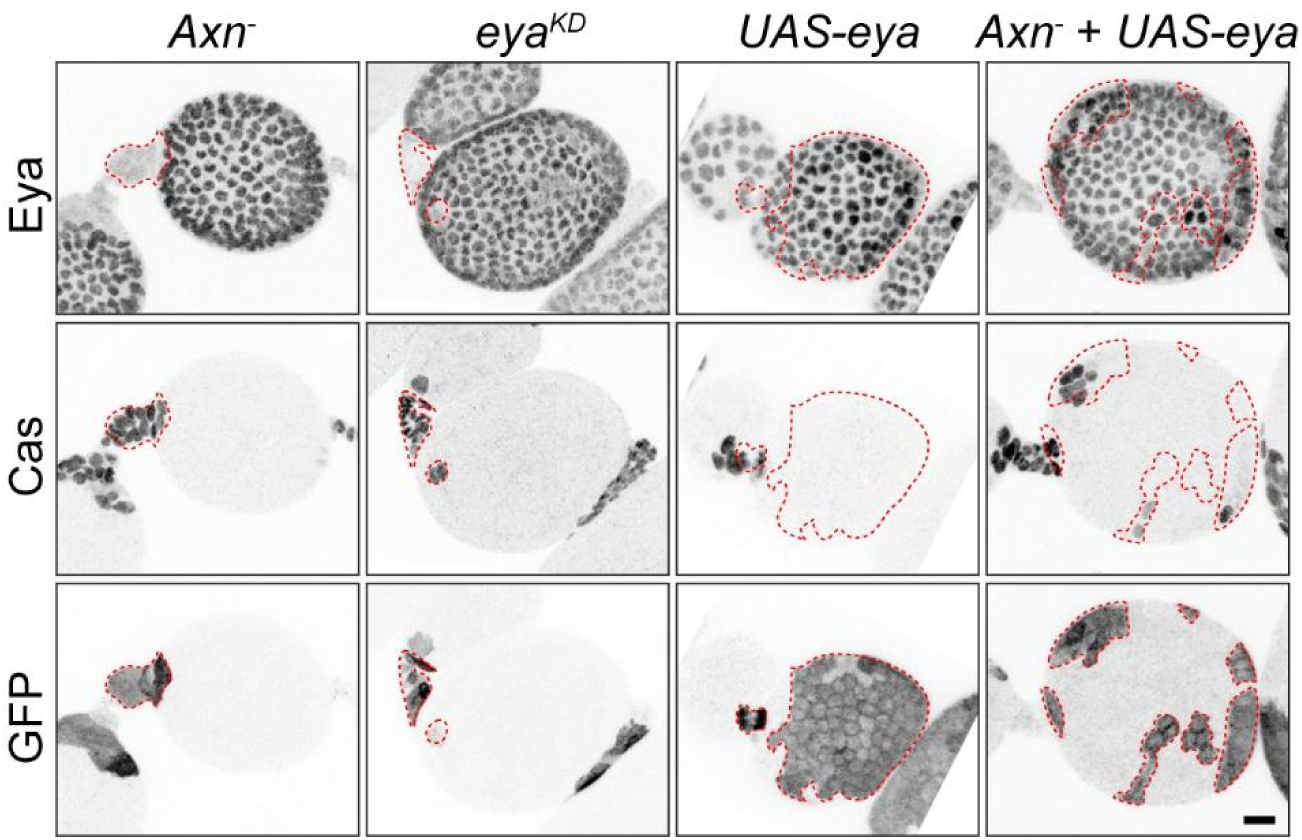
Reduction of Eya is the key cause for the cell fate change in *Axn*^−^. 3D projection view of one half of stage 4 egg chambers with *Axn^S044230^, eyaRNAi*, *UAS-eya*, or *Axn^S044230^* + *UAS-eya* mosaic FSC clones (GFP^+^). Scale bar, 10 μm.

**Supplementary Figure 8.**
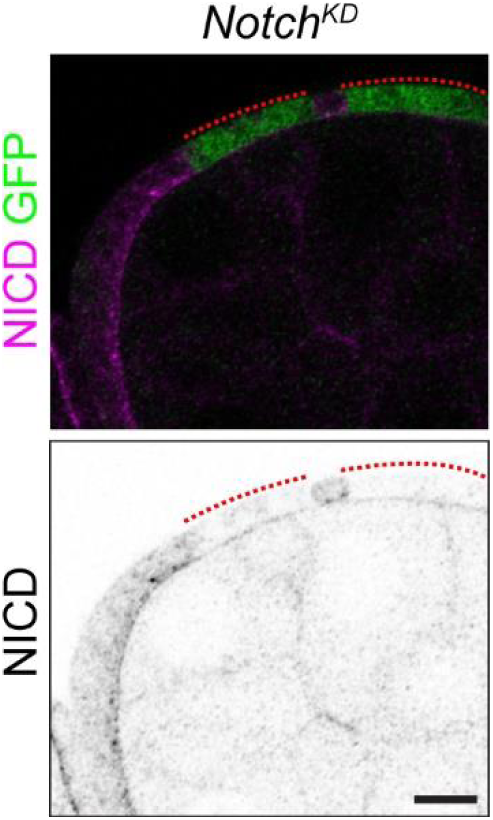
Validation of *notchRNAi*. Sagittal confocal section of *notchRNAi* flip-out clones (GFP^+^) in a stage 6 egg chamber. Scale bar, 10 μm.

**Supplementary Figure 9.**
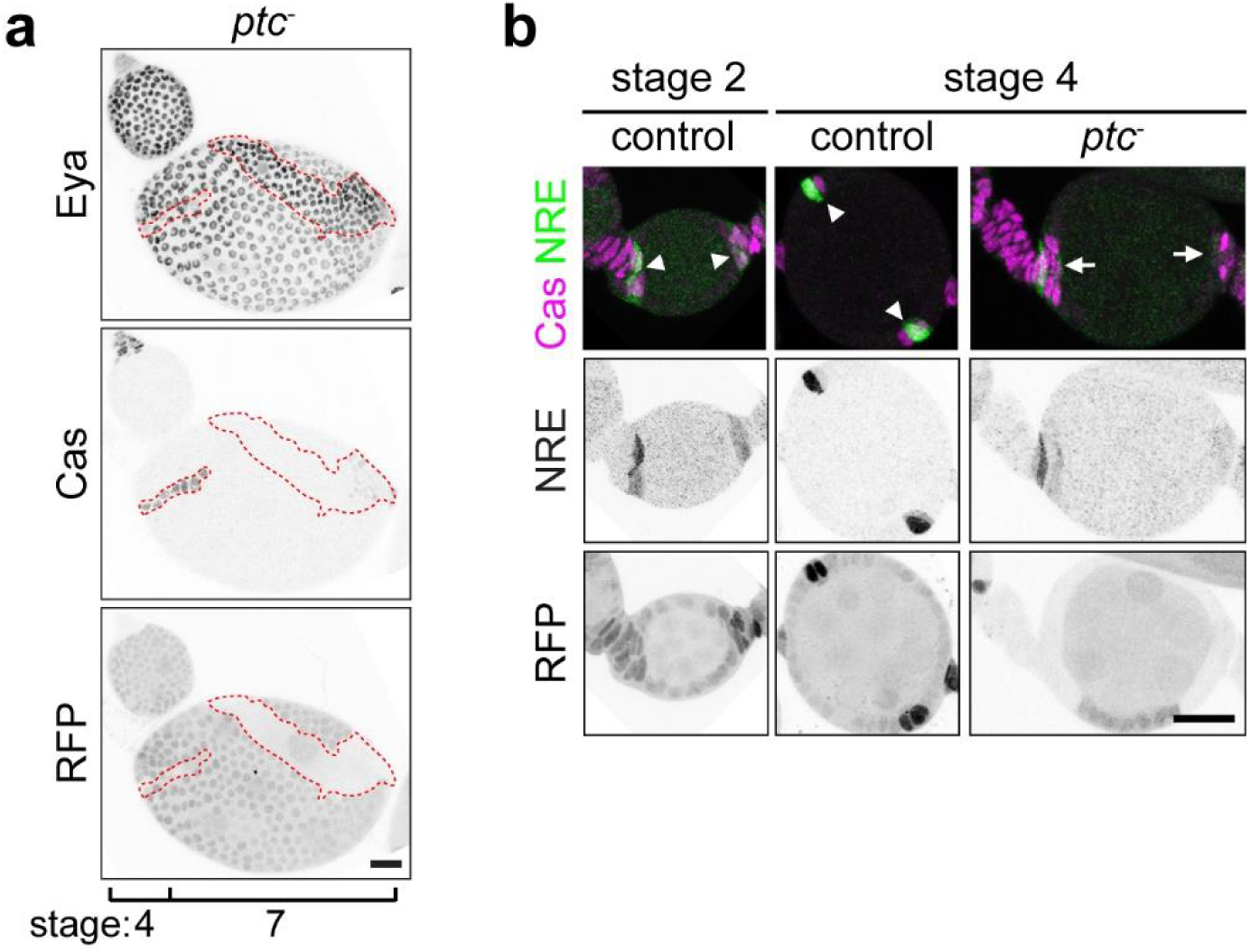
Hh hyper-activation delays follicle cell differentiation. (**a**) 3D projection view of the follicle cell layer in *ptc^S2^* clones (RFP^−^). in a stage 7 egg chamber. (**b**) Notch activity shown by NRE-GFP in *ptc^S2^* heterozygous control (arrowheads) or homozygous mutant (arrows) polar cell regions. Scale bars, 20 μm.

**Supplementary Figure 10.**
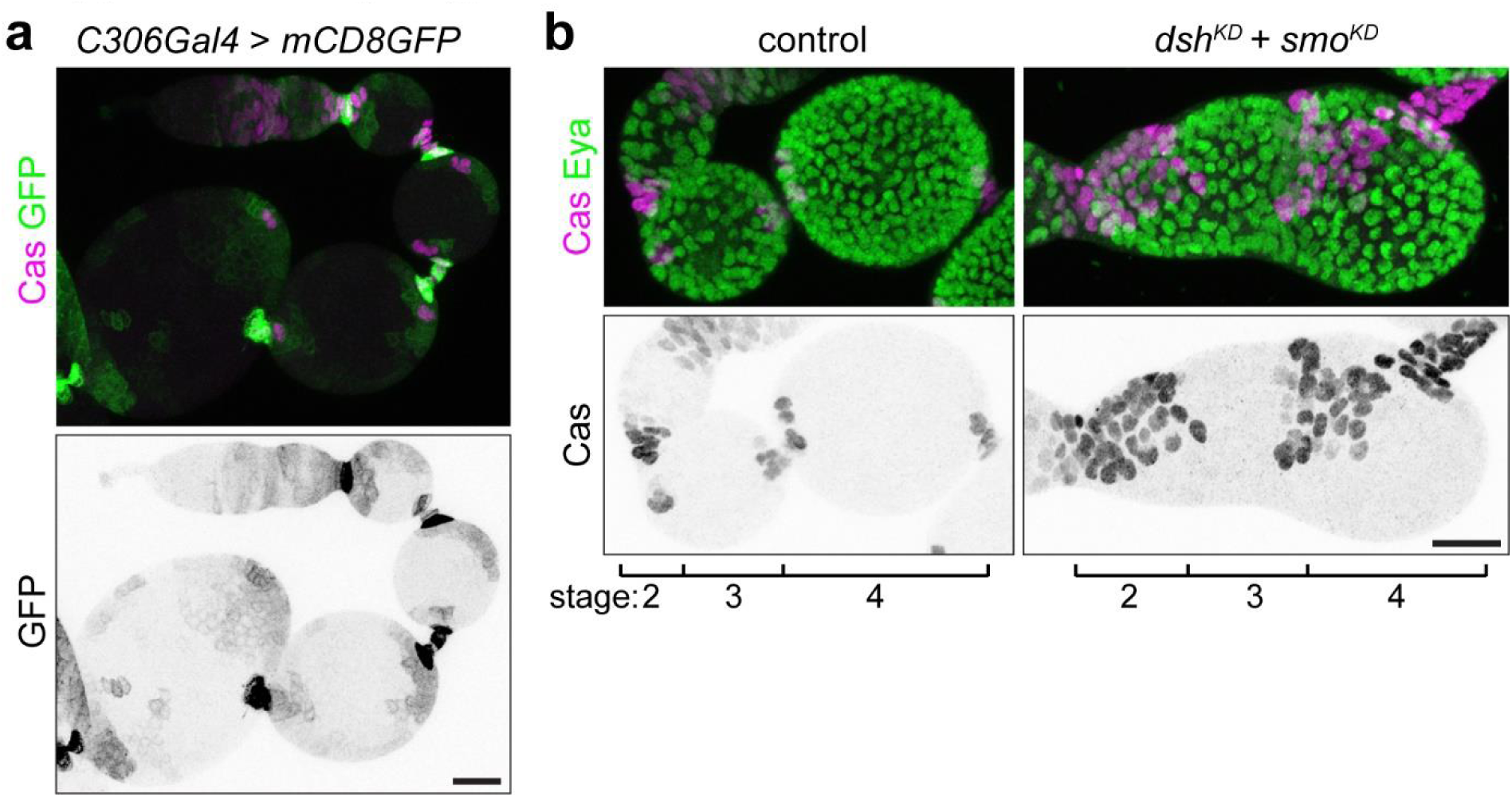
C306 driven knockdown in follicle precursor cells. (**a**) 3D projection view of an ovariole expressing *C306-Gal4* > mCD8GFP. (**b**) 3D projection view of ovarioles with stage 2, 3, 4 egg chambers in *C306-Gal4* > mCD8GFP control or *C306-Gal4* > *dshRNAi* + *smoRNAi*. Scale bars, 20 μm.

**Supplementary Figure 11.**
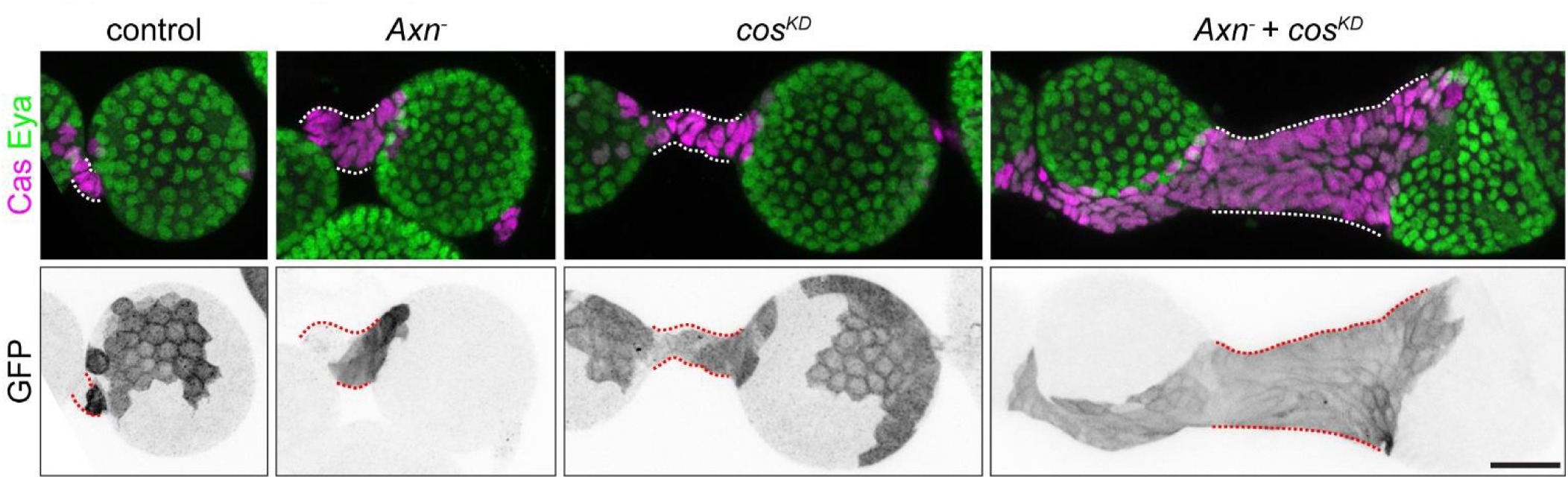
Supernumerary stalk caused by *Axn* and *cos* mutant. 3D projection view of one half of stage 4 egg chambers with FRT82B control, *Axn^S044230^, cosRNAi*, or *Axn^S044230^* + *cosRNAi* mosaic FSC clones. Scale bar, 20 μm.

